# Role of Y box-binding protein 1 (Ybx1/*ybx1*) in zebrafish folliculogenesis: Promoting follicle cell proliferation via suppression of cell cycle inhibitor p21 (*cdkn1a*)

**DOI:** 10.1101/2024.03.27.587099

**Authors:** Bo Zhu, Zhiwei Zhang, Lakhansing Pardeshi, Yingying Chen, Wei Ge

## Abstract

Y box-binding protein 1 (YB-1; Ybx1/*ybx1*) regulates transcription and translation of targeted genes through DNA/RNA-binding. Our research in zebrafish has revealed a high abundance of Ybx1 in the primary growth (PG) follicles in the ovary, which decreases precipitously as the follicles enter the secondary growth (SG) phase. To understand the function of Ybx1 in folliculogenesis, we created an *ybx1* mutant using TALEN and observed a disruption in folliculogenesis in the mutant (*ybx1*-/-) during the transition from previtellogenic (PV) to early vitellogenic (EV) stage of the SG phase, resulting in underdeveloped ovaries and reduced female fertility. Transcriptome and Western blot analyses identified several differentially expressed genes between mutant (*ybx1*-/-) and control (*ybx1*+/-) ovaries. Notably, the expression of *cdkn1a* (p21), a cell cycle inhibitor, increased dramatically in *ybx1*-/- follicles. Disrupting *cdkn1a* gene with CRISPR/Cas9 resulted in embryonic lethality. In p21 heterozygote (*cdkn1a+/-*), however, follicle activation and maturation in the ovary were both advanced, contrasting with the *ybx1-/-* mutant. Interestingly, partial loss of p21 could alleviate the phenotype of *ybx1*-/-. Folliculogenesis resumed in *ybx1*-/-*;p21*+/- females with normal follicle activation (PG-PV transition) and vitellogenic growth (PV-EV transition). Interestingly, the follicle cells from the *ybx1*-/- mutant displayed a poor proliferative activity both in vivo and in vitro; however, the cells from the *ybx1*-/-*;p21*+/- follicles resumed normal proliferation. In conclusion, our study suggests that Ybx1 serves a pivotal role in controlling early folliculogenesis in zebrafish, and its acts, at least partly, by repressing the expression of *cdkn1a,* a cell cycle inhibitor.

## Introduction

Y-box binding protein 1 (YB-1) belongs to the Y-box binding protein family, which also includes YB-2 and YB-3 (1, 2). It consists of a alanine/proline-rich N-terminal domain (A/P domain), a highly conserved cold-shock domain (CSD), and a C-terminal domain (CTD) (3). The CSD is the most conserved nucleic acid-binding domain, which is present in both prokaryotes and eukaryotes. The C-terminal domain may function as a charged zipper to facilitate dimer formation, and it has a strong affinity for single-stranded DNA/RNA in vitro (4). As a DNA/RNA binding protein, YB-1 plays multiple roles in cell functions including DNA replication and repair, transcription, pre-mRNA splicing, mRNA translation, and cell proliferation (1, 5). In cells, YB-1 performs its functions in both cytoplasm and nucleus as RNA binding protein and transcription factor, respectively (6). It may also be secreted from cells to act as a receptor ligand to activate intracellular signal transduction pathways (7). In the mouse, deletion of YB-1 gene (MSY1/*Msy1*) led to late embryonic lethality, indicating importance of YB-1/*Msy1* for early development (8, 9). In zebrafish, deficiency of YB-1 homologue (Ybx1/*ybx1*) induced early maternal embryonic lethality due to gastrulation failure (10) or early embryonic defects (11), suggesting essential roles for YB-1 in zebrafish embryogenesis as well. However, deletion of Y-box-binding protein homologues (CEYs) in *Caenorhabditis elegans* did not induce lethality in the worm (12). Likewise, disruption of Y-box binding proteins (Yps) in *Drosophila* did not affect vitality either (13). These studies suggest functional discrepancies for YB-1 between vertebrates and invertebrates.

In contrast to the ubiquitous expression of YB-1 (14), YB-2 (MSY2/*Msy2*) is germ cell-specific and it regulates transcription and translation specifically in germ cells (15). In mice, both MSY2a and MSY2b are expressed in germ cells (16, 17). MSY2 constitutes 2% of total oocyte proteins and is present in oocytes from diplotene to full-grown stage. It is characterized as an mRNA binding protein to stabilize and store maternal and paternal mRNAs in the cytoplasm (18, 19). The loss of MSY2 in mice resulted in infertility in both males and females with disrupted spermatogenesis and oogenesis (18, 20). In females, MSY2 phosphorylation mediated by CDC2A is associated with oocyte maturation (21). Deletion of MSY2 resulted in progressive loss of oocytes (22), and its knockdown with RNAi reduced fertility due to alteration of protein synthesis and mRNA stability during oogenesis (23). In humans, single nucleotide polymorphisms of MSY2 gene might contribute to susceptibility to spermatogenic impairment in idiopathic infertile men (24). Although YB-2 is present in a wide range of vertebrates including humans and amphibians (25), there has been no loss-of-function studies on YB-2 in species other than rodents.

In contrast to mice and other vertebrates, zebrafish has only one Y-box binding protein, which is YB-1 (Ybx1/*ybx1*) based on sequence information, and there is no homologue for YB-2 in the genome (26). Our recent study showed that despite its similarity to mammalian YB-1 in terms of sequence and ubiquitous expression patterns, zebrafish Ybx1 protein was predominantly present in the gonads (26), which is surprisingly similar to that of YB-2 in other vertebrates. The question then is what is the primary function of Ybx1 in zebrafish? Is it more like YB-1 (MSY1) or YB-2 (MSY2) or both? Also interestingly, Ybx1 protein in the zebrafish ovary is primarily and abundantly present in the follicles of primary growth (PG) stage before follicle activation; however, its level dropped precipitously when the PG follicles were activated to enter the fast-growing SG phase, starting from previtellogenic (PV) stage, and disappeared afterwards in the follicles of vitellogenic stages (26), strongly suggesting a critical role for Ybx1 in controlling early folliculogenesis.

To address the above questions, we performed a loss-of-function study on Ybx1 in zebrafish by transcription activator-like effector nuclease (TALEN) technology followed by phenotype analysis with particular emphasis on reproduction, especially ovarian folliculogenesis. As expected, the loss of Ybx1 resulted in severe post-hatching lethality, which was similar to that of YB-1 in mice. Interestingly and fortunately, the mutant did not show complete penetrance as some individuals managed to survive to adulthood, allowing for studies on reproductive development and functions. Similar to that of YB-2 (MSY2) in mice, the mutant zebrafish females (*ybx1-/-*) showed severe defects in ovarian folliculogenesis. The follicles were mostly arrested at the PV stage with normal formation of cortical alveoli but not yolk granules, and subsequent development to advanced vitellogenic stages was largely blocked. Further experiments using CRISPR/Cas9 method demonstrated that the role of Ybx1 in early follicle development involved controlling follicle cell proliferation through p21 (*cdkn1a*), a cell cycle inhibitor.

## Materials and methods

### Animals

The wild type (WT) zebrafish (*Danio rerio*) of AB strain was used in this study. The fish were maintained in the ZebTEC multilinking rack system (Tecniplast; Buguggiate, Italy) under an artificial photoperiod of 14-h light:10-h dark. The temperature, pH, and conductivity of the system were 28 ±1C°, 7.5, and and 400 µS/cm, respectively. The fish were fed twice a day with Otohime fish diet (Marubeni Nisshin Feed, Tokyo, Japan) by the Tritone automatic feeding system (Tecniplast).

All experiments were performed under a license from the Government of the Macau Special Administrative Region (SAR), and the protocols were approved by the Animal Ethics Panel of the University of Macau (Approval No. AEC-13-002).

### Mutagenesis of ybx1 and cdkn1a genes

To disrupt *ybx1* gene, the sequence of *ybx1* gene was retrieved from the Ensembl database (ENSDARG00000004757) for target site identification. The sequence in the first exon near the ATG start codon was chosen for targeting by TALEN, and both the left and right TALEs were designed by the online software (TAL Effector Nucleotide Targeter 2.0 Tools) (https://tale-nt.cac.cornell.edu/node /add/talen). The target sequences are as follows: TGCCGCCGACGCGGAGA (left TALE), GCCGCAGCTACCGCGGGGGA (right TALE), and GCCCGTCCAGCCCGGCA (spacer sequence) (Fig. S1A). The left and right TALE mRNAs were transcribed by the Ambion SP6 mMessage Machine Kit (Thermo Fisher Scientific, Waltham, MA), quantified by the NanoDrop 2000c (Thermo Fisher Scientific), and co-injected into 1-2 cell embryos (200 pg each) to generate mosaic F0 mutants. The F0 founders carrying mosaic mutations were identified by genotyping on genomic DNA extracted from the caudal fin, and they were crossed with a WT fish to obtain F1 generation. The F1 individuals carrying mutations were identified and characterized by TA cloning and sequencing. A mutant line carrying a 17-bp deletion was selected for lineage establishment and phenotype analysis.

For *cdkn1a* (p21) mutagenesis, we used CRISPR/Cas9 method instead of TALEN. Similarly, the gene sequence of *cdkn1a* was obtained from the Ensembl database (ENSDARG00000076554) for target site identification. The region in the first exon near the ATG start codon was selected for targeting (Fig. S2A). The targeting sgRNA was designed using the online software ZiFiT Targeter (Vision 4.2) (http://zifit.partners.org/ZiFiT/ChoiceMenu.aspx). The oligos for *cdkn1a* sgRNA synthesis were TAGGGCGGAGACTTCCACTGGG and AAACCCCAGTGGA AGTCTCCGC. The sgRNA and Cas9 mRNA (100 pg each) were co-injected into 1-2 cell embryos to generate F0 generation. The F0 fish carrying mutations were crossed with the WT fish to generate F1 generation. A mutation with 14 bp deletion was selected for lineage establishment and phenotype analysis.

### Genotyping

We used two methods in this study for genotyping individuals: high-resolution melt analysis (HRMA) and heteroduplex mobility assa*y* (HMA). For HRMA assay, real-time PCR was carried out on the CFX96 Real-Time PCR Detection System (Bio-Rad Laboratories, Hercules, CA) in white PCR plates (96 wells) to generate 90-150 bp amplicons. The reaction was run in 10 µl mixture containing genomic DNA, two primers and 1× SsoFast EvaGreen Supermix (Bio-Rad). Three-step PCR was carried out: 95°C for 3 min and 35 cycles of 95°C for 20 sec, 62°C for 20 sec and 72°C for 20 sec. Melting curves of PCR amplicons were obtained with temperatures ranging from 65°C to 95°C. Data acquisition was performed for every 0.2°C increase in temperature, with 10 sec per step (Fig. S1B and S2C). For HMA analysis, the PCR products from HRMA were electrophoresed on 20% polyacrylamide gels at 30 V for at least 10 h, and the gel was stained with 0.01% GelRed dye (Biotium, Hayward, CA) for 10 min and visualized on the Gel Doc XR System (Bio-Rad) (Fig. S1C).

The deletion of target genes was also validated by PCR on genomic DNA with a mutant-specific primer with 3’-end overhanging the indel sequence. The PCR reaction (10 μL) was electrophoresed on 1% agarose gel, stained with GelRed dye, and visualized on the Gel Doc XR System (Bio-Rad). Signals could be amplified from control DNA samples but not the mutants (Fig. S1D).

### Western blot analysis

For immunoblotting, the ovarian tissues or follicle cells from control and mutant females were lysed in 100 μL 1x SDS sample buffer (62.5 mM Tris-HCl, 1% w/v SDS, 10% glycerol, 5% mercaptoethanol, pH=6.8). The lysates were heated at 95°C for 10 min and centrifuged at 12,000 rpm for 15 min at 4°C, followed by determination of protein concentrations. The samples (total 100 μg protein from each fish) were separated on 12% polyacrylamide gels and transferred to PVDF membranes. The membranes were blocked with 5% non-fat milk in 1x TBST for 1 h at room temperature. After washing three times with 1x TBST, the membranes were incubated overnight at 4°C with anti-zebrafish Ybx1 primary antibody we prepared previously (1:2000) (GenScript, Nanjing, China) (26), anti-p21 (1:2000) (C-19, Santa Cruz Biotechnology, CA) and anti-β-actin (1:2000) (13E5 rabbit mAb, Cell Signaling Technology, MA) in 10 ml of 1x TBST with 5% non-fat milk. They were washed three times with 1x TBST followed by incubation with the HRP-conjugated secondary antibody (1:2000) (Cell Signaling Technology) in 10 ml 1x TBST for 1 h at room temperature. After washing, the membranes were incubated with the SuperSignal West Femto Maximum Sensitivity Substrate (Thermo Fisher Scientific) and signals detected on the ChemiDoc MP imaging system (Bio-Rad).

### Immunohistochemistry (IHC)

The zebrafish ovaries from the controls (+/+ and/or +/-) and mutants (-/-) were dissected and fixed in 1 mL Bouin’s solution overnight at room temperature. After embedding in paraffin, the ovaries were sectioned at 7 μm and mounted on Superfrost Plus slides (Thermo Fisher Scientific). The slides were deparaffinized and rehydrated followed by antigen retrieval for 10 min in sodium citrate buffer (10 mM) at sub-boiling temperature (95°C). The endogenous hydrogen peroxidase was inactivated by treating the sections with 3% H_2_O_2_ for 10 min. The sections were then washed with 1x PBS three times at 5 min interval before blocking for 1 h with normal horse serum at room temperature. Slides were incubated overnight at 4°C with 100 μL Ybx1 and p21 primary antibodies in blocking solution with normal horse serum (1:2000 dilution for both), washed with 1x PBS three times, and then incubated with 100 μL secondary antibody for 1 h at room temperature. After incubation with the avidin/biotinylated enzyme complex (ABC) from the VECTASTAIN ABC-HRP Kit (Vector Laboratories, Burlingame, CA) for 30 min at room temperature, the slides were stained with 3,3’-diaminbenzidine (DAB) (Vector Laboratories) for 5 min and the staining stopped in running tap water. The slides were then dehydrated and mounted for observing and photographing on the Nikon ECLIPSE Ni-U microscope (Nikon, Tokyo, Japan).

### Immunofluorescence (IF)

Ovarian tissues were fixed in 4% paraformaldehyde (PFA) solution prior to paraffin sectioning. After antigen retrieval and blocking with serum, the sections were incubated first with anti-Ybx1 primary antibody and then Alexa Fluor 488/568-conjugated secondary antibody (Life Technologies, Carlsbad, CA). The sections were counterstained for nuclei with Hoecyst33342 ((Thermo Fisher Scientific) before observation and photographing on the ECLIPSE Ni-U microscope (Nikon).

### Histological examination

The fish of different genotypes (*+/+*, *+/-* and *-/-*) were photographed using a digital camera (EOS700D, Canon; Tokyo, Japan) to analyze gross morphology. Morphometric parameters were measured, including standard body length (BL), body weight (BW), and gonadosomatic index (GSI; gonadal weight/body weight). For histological examination, ovarian samples from different genotypes including single and double mutants and the whole body of juvenile fish were immediately fixed in Bouin’s solution overnight at room temperature. The fixed samples were dehydrated and embedded in paraffin according to our previous reports (27, 28), and then serially cut into 7 μm sections on a microtome (Leica, Wetzlar, Germany). Slides were stained with hematoxylin and eosin (H&E), then observed and photographed on the Nikon ECLIPSE Ni-U microscope (Nikon).

### Fecundity assay

Four female individuals were randomly selected for fecundity assay for each genotype (*ybx1+/+*, *ybx1+/-* and *ybx1-/-*; *cdkn1a+/+* and *cdkn1a+/-*). Every female of each genotype was bred with two WT males in a breeding tank. The breeding assay was conducted three times for each individual female tested, with 3 or 4-day interval between the assays. Embryos were collected and counted after spawning. Non-fertilized embryos were removed after 24 hours post-fertilization (hpf) to calculate fertilization rate.

### Follicle isolation

Zebrafish were anesthetized by cold shock on ice and decapitated before dissection. Ovaries were carefully removed from three female zebrafish and placed in a 100-mm Petri dish containing 60% Leibovitz L-15 medium (Invitrogen, Waltham, MA). Fat tissues and ligaments surrounding the ovaries were stripped off using a BD Microlance 26G needle (BD, San Diego, CA). To isolate PG and PV follicles from the ovary, a transfer pipette (JET Biofil, Guangzhou, China) was used to pipet ovarian fragments up and down a few times, followed by further pipetting with a BD 23G needle for a few times. After separation of follicles, the PV follicles were isolated by filtering through sieves with pore sizes of 180 and 250 μm (JiuFeng, Heng Shui, China). The PV follicles retained between 180 and 250 μm were collected by centrifugation. Similarly, the PG follicles (< 100 μm) from each genotype were isolated by filtering through sieves with 100-μm pore size (SPL Lifesciences, Waunakee, WI). The PG follicles that passed through the 100 μm sieve were collected by centrifugation at 3000 *rpm* for 2 min. The isolated PG and PV follicles were washed twice with 1 x PBS. In total, nine zebrafish were used for each genotype and divided into three groups (3 fish each) for follicle isolation and sampling.

### Transcriptome data analysis

Total RNA from PG follicles (*cdkn1a+/+* and *cdkn1a+/-*) and PV follicles (*ybx1+/+* and *ybx1-/-*) of each group was extracted using Tri-Reagent (Molecular Research Center, Cincinnati, OH) according to the protocol of the manufacturer and our previous report (29). The RNA was then treated with DNase for 10 min at 37^◦^C to remove genomic DNA [10 μg RNA in 100 μl reaction buffer with 2U DNase I from NEB (Ipswich, MA)] followed by phenol-chloroform extraction and ethanol precipitation. For RNA library construction, the integrity of the RNA samples was first analyzed on the Bioanalyser 2100 (Agilent, Stockport, UK). RNA libraries were prepared using the NEBNext Ultra Directional RNA Library Prep Kit (NEB) and sequenced on the HiSeq 2500 Sequencing System (Illumina, San Diego, CA) with 100-bp paired end reads.

The transcriptome data were analyzed at the Genomics and Bioinformatics Core (Faculty of Health Sciences, University of Macau), and all the raw data were submitted to NCBI database with accession numbers GSE239622 and GSE239623. Methods for analyzing differentially expressed genes (DEGs), Gene Ontology (GO) and Kyoto Encyclopedia of Genes and Genomes (KEGG) enriched pathways, were according to our previous report (30).

### Reverse transcription and real-time quantitative PCR (RT-qPCR)

Total RNA was extracted from follicles using Tri-Reagent (Molecular Research Center). Reverse transcription was performed at 37°C for 2 h in a total volume of 10 μl reaction solution containing 3 μg RNA, 0.5 μg oligo (dT), 1X MMLV RT buffer, 0.5 mM each dNTP, 0.1 mM dithiothreitol, and 100 U M-MLV reverse transcriptase (Invitrogen). The expression of several representative genes was determined by RT-qPCR in PG follicles, including *vtg1*-*vtg7*. The expression levels were normalized to that of the housekeeping gene *ef1a*. The standard for each gene in the assay was prepared by PCR amplification of the cDNA fragment with specific primers (Supplemental Table S1). The qPCR was performed on the CFX96 Real-time PCR Detection System (Bio-Rad).

### ELISA assay

To further confirm and measure the expression of Vtg proteins in PV follicles, we used enzyme-linked immunosorbent assay (ELISA) evaluate Vtg protein levels in PV follicles by ELISA (Biosense Laboratories, Bergen, Norway). Briefly, the PV follicles were isolated from the ovaries and used for sample preparation. The ELISA was performed according to the protocol of the Zebrafish Vitellogenin ELISA kit (V01008402, Biosense Laboratories).

### Primary culture of follicle cells

The primary culture of zebrafish ovarian follicle cells was performed according to our previous studies (31–33) with minor modifications. Briefly, the PV follicles were isolated from four different genotypes: *ybx1+/+;p21+/+*, *ybx1+/+;p21+/-*, *ybx1*-/-*;p21*+/+ and *ybx1*-/-*;p21*+/-. Three females were used for each genotype. The isolated PV follicles were placed in 12-well plates (Corning, Brooklyn, NY) after washing with 60% Leibovitz L-15 medium. They were incubated for 6 days at 28°C with 5% CO_2_ in medium M199 supplemented with 10% fetal calf serum (HyClone, Logan, UT). At the end of incubation, the attached PV follicles and the proliferated follicle cells were photographed on the Nikon Eclipse TS100 inverted microscope (Nikon) for measuring the diameters of the cell outgrowth zone.

### MTT assay on follicle cell proliferation

To evaluate follicle cell proliferation in vitro, the follicle cells were harvested on the sixth day of follicle incubation by trypsinization with trypsin-EDTA (0.25%) (Thermo Fisher Scientific). These cells were then sub-cultured into 96-well plates (SPL Life Sciences, Pocheon-si, Korea). On the third day of subculture, a proliferation assay was conducted on the cultured follicle cells using the MTT Cell Proliferation and Cytotoxicity Assay kit (Beyotime, Shanghai, China). Briefly, the MTT solution (5 mg/ml, 10 µL) was added to each well to incubate for 4 h followed by addition of 100 μL formazan solution. The plates were carefully mixed and incubated at 28°C for several hours until purple crystals had dissolved completely. Absorbance was then measured at 570 nm on the Infinite M200 Pro spectrophotometer (Tecan, Männedorf, Switzerland).

### BrdU incorporation assay on follicle cell proliferation

To evaluate follicle cell proliferation in vivo, we performed BrdU (5-bromo-2’-deoxyuridine) incorporation assay. Briefly, the BrdU solution (10 μg/mL, 25 µL) was injected into each fish with a microliter syringe and the injected individuals were maintained at 28°C for 4-6 h. The ovaries were dissected and fixed in Bouin’s solution, embedded in paraffin after dehydration, and then serially cut into 7-µm sections on a microtome (Leica). Slides were immersed in 2N HCl for 15 min for DNA denaturation, rinsed three times with PBS (2 min each), and blocked by 5% normal horse serum for 1 h at room temperature. The sections were then incubated with anti-BrdU antibody (1:200; Sigma Aldrich, St. Louis, Missouri) overnight in a cold room. The slides were then rinsed with PBS three times (2 min each) followed by incubation for 2 h at room temperature with the secondary antibody for IHC staining. The sections were observed and photographed on the Nikon ECLIPSE Ni-U microscope (Nikon).

### Data analysis

Gene expression was assessed by quantifying mRNA levels of target genes via qPCR, with normalization to the internal control, *ef1a*. Quantitative data were analyzed using either one-way ANOVA or Student’s t-test, as appropriate, with Prism software (GraphPad Software, San Diego, CA) running on Apple OS X. Results are expressed as mean ± SEM.

## Results

### Establishment of ybx1-/- mutant zebrafish line

Using TALEN method, we targeted the first exon of *ybx1* gene and established several mutant lines for *ybx1* gene with different indel mutations and we chose the line with 17-bp deletion for phenotype analysis (Fig. S1A). We characterized this mutant line with HRMA, HMA, PCR, and Western blot to confirm the mutation of *ybx1* gene. With HRMA, we could identify three distinct melt curves for *ybx1+/+*, *ybx1+/-* and *ybx1-/-,* respectively (Fig. S1B). The HMA assay displayed three different banding patterns for *ybx1+/+*, *ybx1+/-,* and *ybx1-/-* (Fig. S1C). Using a mutation-specific primer that overhangs the deleted sequence at its 3’-end, we could detect specific PCR signal in both *ybx1+/+* and *ybx1+/-*, but not *ybx1-/-* individuals (Fig. S1D). We also performed Western blot on proteins extracted from the ovary using a specific antibody against zebrafish Ybx1 (26). The result showed the absence of Ybx1 protein in *ybx1-/-,* but not *ybx1+/+* and *ybx1+/-* (Fig. S1E).

The successful deletion of *ybx1* gene was also confirmed by IHC and IF staining, which showed a complete lack of signals for Ybx1 protein in the mutant ovary (*ybx1-/-*) compared with the control (*ybx1*+/+). In the control ovary, Ybx1 protein was abundantly expressed in the PG follicles, primarily in the oocytes. As the follicles transitioned to the PV stage, Ybx1 level diminished markedly and became undetectable after the onset of vitellogenic growth, which is characterized by the accumulation of yolk granules (Fig. 1A-E), in agreement with our recent report (26). Although oocyte is the primary site where Ybx1 is produced, we also observed strong IHC signal in the follicle cells (Fig. 1F). All these lines of evidence indicate clearly that the *ybx1* gene was successfully deleted from the zebrafish genome.

**Fig 1.**
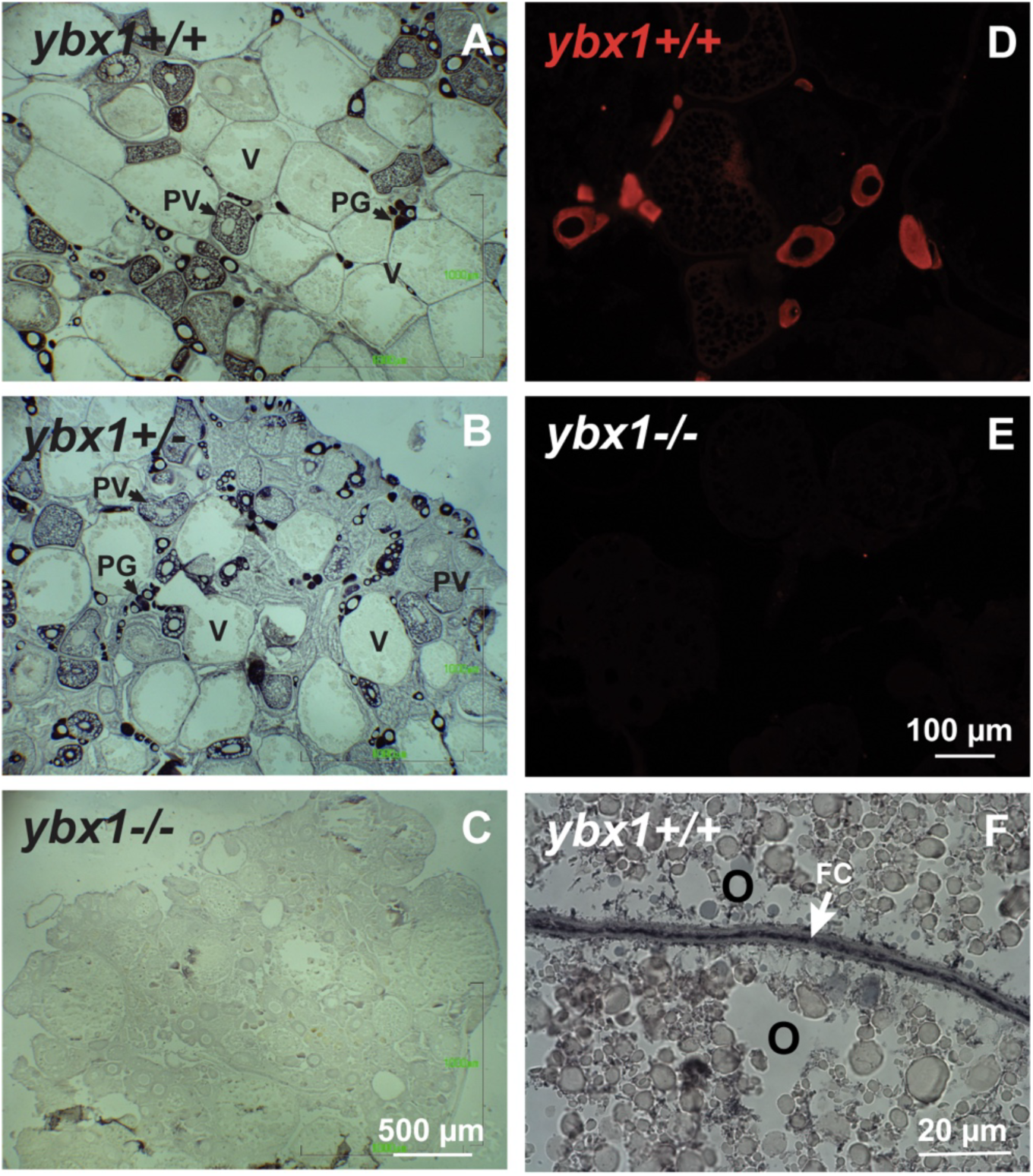
Immunohistochemical (IHC) and immunofluorescent (IF) staining for Ybx1 in the ovaries of controls and mutant (*ybx1*-/-). (A-C) IHC staining for Ybx1 in the ovaries of controls (*ybx1+/+* and *ybx1+/-*) and mutant (*ybx1*-/-). No Ybx1 was detected in the mutant ovary. In the control ovaries, the expression of Ybx1 protein was the highest in the PG follicles, and its signal started to decline at PV stage. PG, primary growth; PV, previtellogenic; V, vitellogenic. (D and E) IF staining for Ybx1 in the control (*ybx1+/+*) and mutant (*ybx1-/-*) ovaries. (F) IF staining for Ybx1 in the follicle cells (FC). O, oocyte.

### Ovarian atrophy in ybx1 mutant

Ybx1 plays a vital role in zebrafish development and survival as the loss of Ybx1 in *ybx1-/-* resulted in a significant post-hatching mortality with only 7% survival rate (data not shown). Interestingly, the survived individuals could grow normally to the adult stage without significant difference from the controls in terms of body size (BW and BL) (Fig. 2A). This fortunately allows for investigating the roles of Ybx1 in controlling ovarian development and folliculogenesis in sexually mature adults. Morphological examination revealed that the loss of *ybx1* resulted in severe ovarian atrophy as shown by the significantly reduced GSI (180 dpf) (Fig. 2B). The ovarian atrophy of the mutant (*ybx1*-/-) was apparent as early as 60 dpf and became increasingly pronounced at later stages (90-180 dpf), coinciding with the transition of control ovaries into the fast-growing vitellogenic phase (Fig. 2C), indicating a crucial role for Ybx1 in ovarian growth.

**Fig 2.**
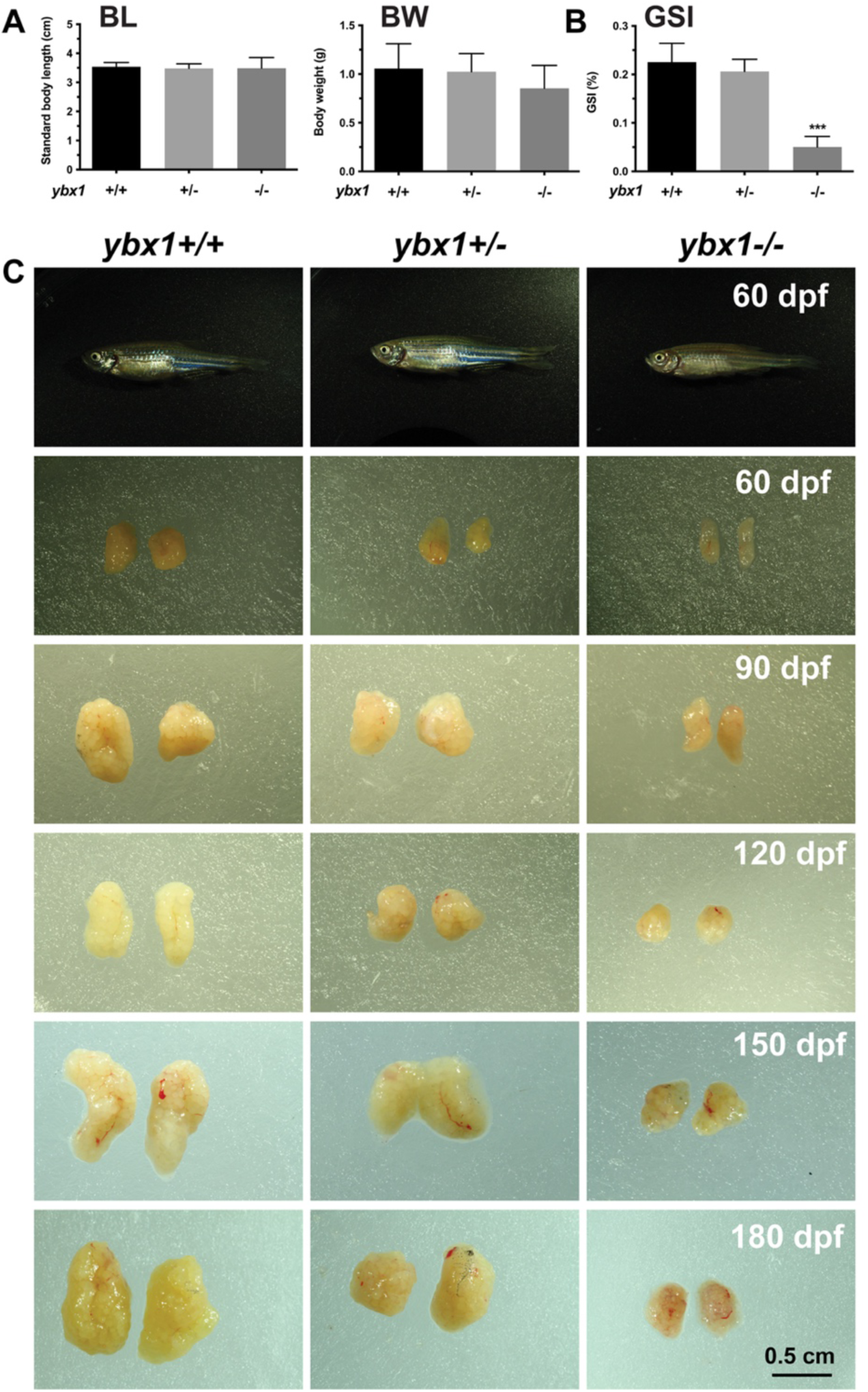
Effects of *ybx1* mutation on somatic growth and ovarian development. (A) Effects of *ybx1* mutation on body growth and reproduction. BL, standard body length; BW, body weight. (B) Gonadosomatic index (GSI) of female controls (*ybx1+/+* and *ybx1+/-*) and mutant (*ybx1*-/-). The mutant females exhibited significantly lower GSI compared to the controls. (C) Ovarian growth in controls (*ybx1+/+* and *ybx1+/-*) and mutant (*ybx1*-/-). The mutant ovary showed significant atrophy or hypotrophy from 60 to 180 dpf.

### No effect of ybx1 mutation on follicle activation or PG-PV transition

As reported recently (26) and observed in this study, Ybx1 protein was abundant in PG follicles and its level declined after the follicles entered the PV stage, suggesting a potential role for Ybx1 in PG-PV transition. The PG-PV transition marks follicle activation in folliculogenesis and puberty onset when it occurs for the first time in the developing ovary (34). To test if Ybx1 plays any roles in follicle activation, we first examined early follicle development at puberty onset, in particular the transition from PG to PV stage when the oocyte starts to accumulate cortical alveoli. Unexpectedly, the lack of Ybx1 did not cause any change in PG-PV transition as PV follicles with cortical alveoli could form well in *ybx1-/-* mutant ovary around 45 dpf, fully comparable to those of the age-matched controls (*ybx1+/+* and *ybx1+/-*). IHC staining confirmed the absence of Ybx1 protein in the mutant (Fig. 3).

**Fig 3.**
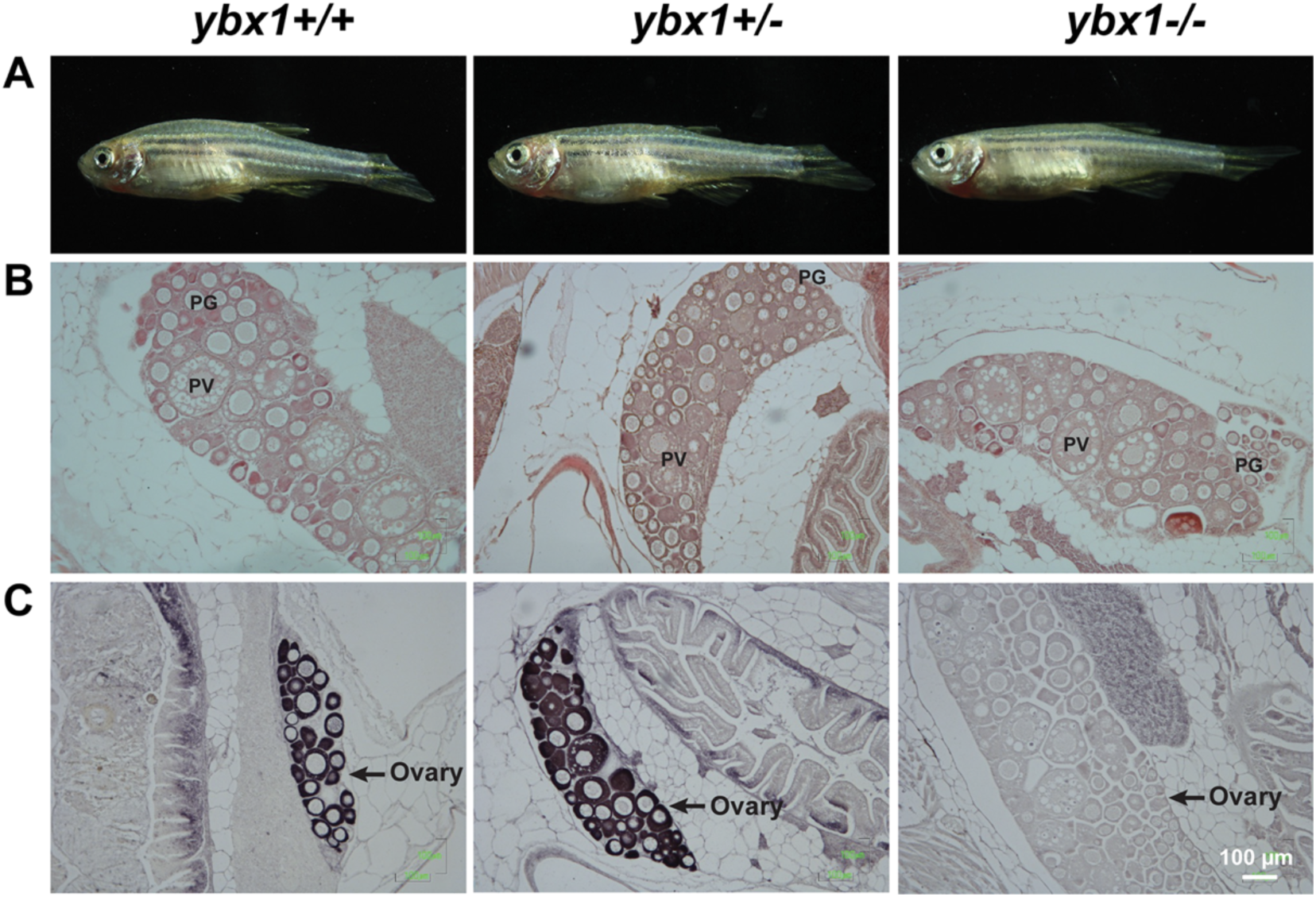
Analysis of follicle activation (PG-PV transition) in juvenile *ybx1* mutant. (A) Female fish of three genotypes at 41 dpf: controls (*ybx1*+/+, *ybx1*+/-) and mutant (*ybx1*-/-). (B) H&E staining of ovarian sections. All three genotypes possessed both PG and PV follicles in the ovaries, indicative of a normal PG-PV transition or follicle activation. PG, primary growth; PV, pre-vitellogenic. (C) IHC staining for Ybx1 protein in the ovaries. The follicles (mostly PG) were strongly stained for Ybx1 in control ovaries (*ybx1*+/+, *ybx1*+/-), whereas no Ybx1 staining was detected in the mutant (*ybx1*-/-) ovary.

### Blockade of follicle development at PV-EV transition in ybx1-/- ovary

Having demonstrated no effect of *ybx1* mutation on PG-PV transition, we went on to examine folliculogenesis in adults (Fig. 4A). Morphometric analysis at 180 dpf showed that the ovaries in the control fish (*ybx1+/+* and *ybx1+/-*) developed normally, in sharp contrast to the underdeveloped (hypotrophic) ovaries of the mutant (*ybx1-/-*) (Fig. 4B). Histological examination showed all stages of follicles in the control ovaries (*ybx1+/+* and *ybx1+/-*) from PG to full-grown stage (FG); however, in the mutant ovary, the follicles were arrested at PV stage with normal formation of cortical alveoli but without accumulation of yolk granules, indicating lack of vitellogenesis with follicles blocked at the transition from PV to early vitellogenic stage (EV) (Fig. 4C and D).

**Fig 4.**
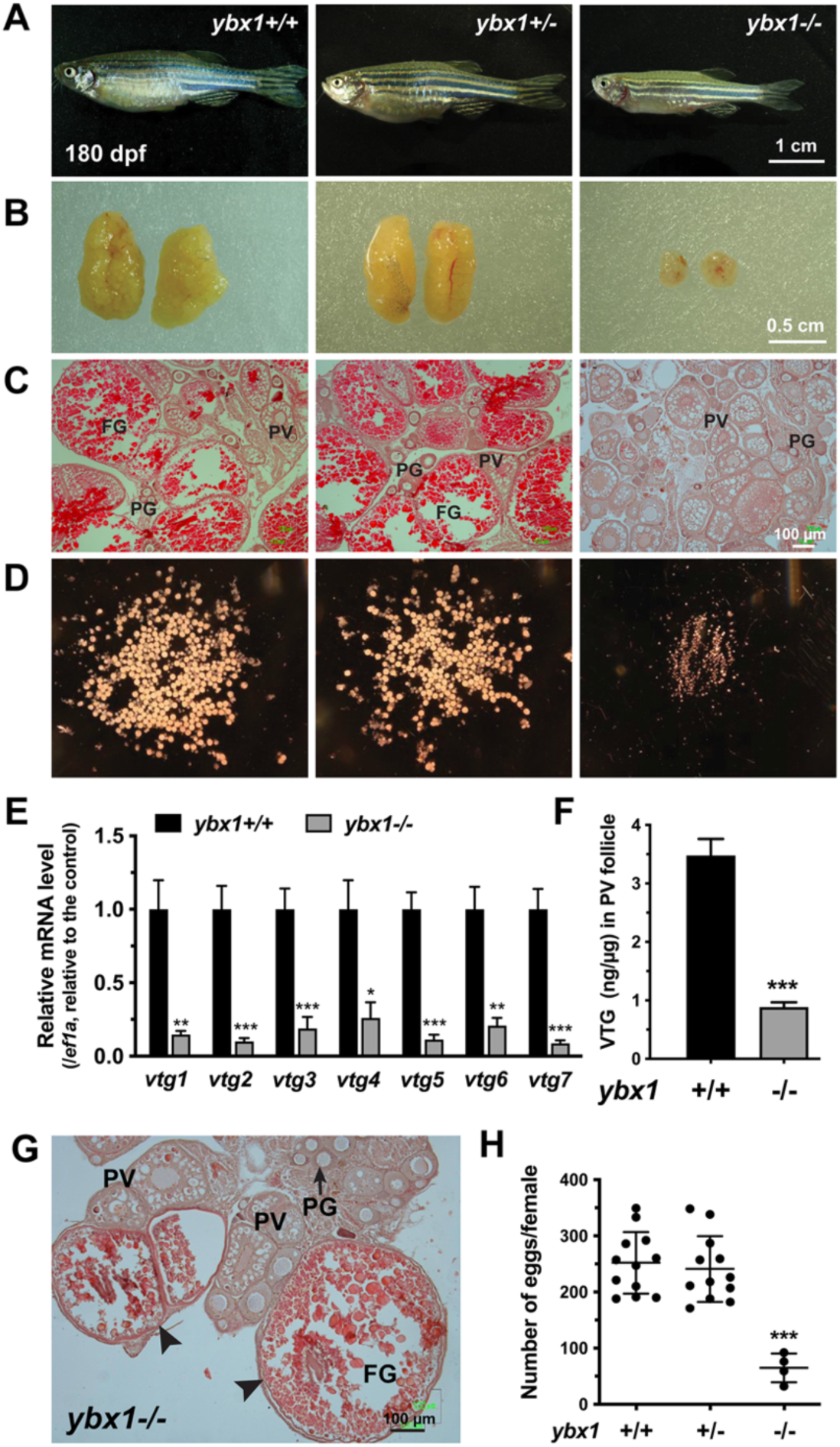
Phenotype analyses of adult *ybx1* mutant. (A) Gross morphology of controls (*ybx1*+/+, *ybx1*+/-) and mutant (*ybx1*-/-) at 180 dpf. (B) Gross anatomy of ovaries from different genotypes. (C) Histology of the control and mutant ovaries. (D) Dispersed follicles from the controls and mutant. (E) Expression of vitellogenin genes (*vtg1-7*) in the PV follicles from WT control ((*ybx1*+/+) and mutant (*ybx1*-/-). (F) Vitellogenin (VTG) levels in the WT control and mutant PV follicles. (G) Vitellogenic follicles in the mutant ovary (*ybx1*-/-). (H) Fecundity (number of eggs spawned by each female) of controls (*ybx1*+/+, *ybx1*+/-) and mutant (*ybx1*-/-). PG, primary growth; PV, pre-vitellogenic; FG, full-grown; * P < 0.05, ** P < 0.01 and *** P < 0.001 compared with the control group.

The phenotype of *ybx1-/-* was surprisingly similar to that of bone morphogenetic protein 15 (*bmp15*) mutant whose follicle development was also blocked at the PV stage without entering vitellogenic growth (35). On interesting observation in the *bmp15*-/- ovary was the reduced expression of all vitellogenin genes (*vtg1-7*) in the mutant follicles (35). This prompted us to analyze the expression of these genes in the PV follicles of the *ybx1*-/- mutant. Surprisingly, all seven Vtg genes (*vtg1-7*) significantly reduced their expression in the mutant PV follicles (Fig. 4E). Consistently, the level of Vtg protein also decreased dramatically in the mutant PV follicles (Fig. 4F).

Interestingly, we discovered by histology that a few follicles could eventually overcome the blockade to enter the vitellogenic growth at later stages of development (>180 dpf) (Fig. 4G), resulting in female sub-fertility. Fertility test showed that out of 68 breeding tests on mutant females, only 4 tests yielded fertilized eggs (4/68), much lower than the success rates in the controls (*ybx1+/+*: 12/16; *ybx1+/-*: 12/15). The number of eggs obtained from the 4 mutant females was less than 100, also significantly lower than those from the control females (>200/fish) (Fig. 4H).

### In vitro and in vivo evidence for impaired follicle cell proliferation in ybx1-/- ovary

We have previously reported that zebrafish follicles can be cultured in vitro, and during the 6-day incubation period, the follicle cells can actively proliferate, radiating outward from the attached oocyte to form an outgrowth or halo-like ring of cells. Interestingly, the follicle cells from different stages of folliculogenesis exhibited varying proliferative capabilities. The PG follicles prior to the fast-growing secondary growth stage or vitellogenic growth and the post-vitellogenic FG follicles displayed the lowest proliferation activity (32). This in vitro follicle culture serves as an excellent assay to evaluate the proliferative ability of follicle cells. Using this approach, we cultured PV follicles from both the control (*ybx1*+/+) and mutant (*ybx1*-/-) fish in vitro. Surprisingly, at the end of the 6-day incubation, the control and mutant follicles exhibited a significant difference in follicle cell proliferation. A large ring of cell outgrowth formed around each control follicle, while in contrast, very few follicle cells grew from the mutant follicle (Fig. 5A), indicating a severe impairment of follicle cell proliferation in *ybx1-/-* mutant.

**Fig 5.**
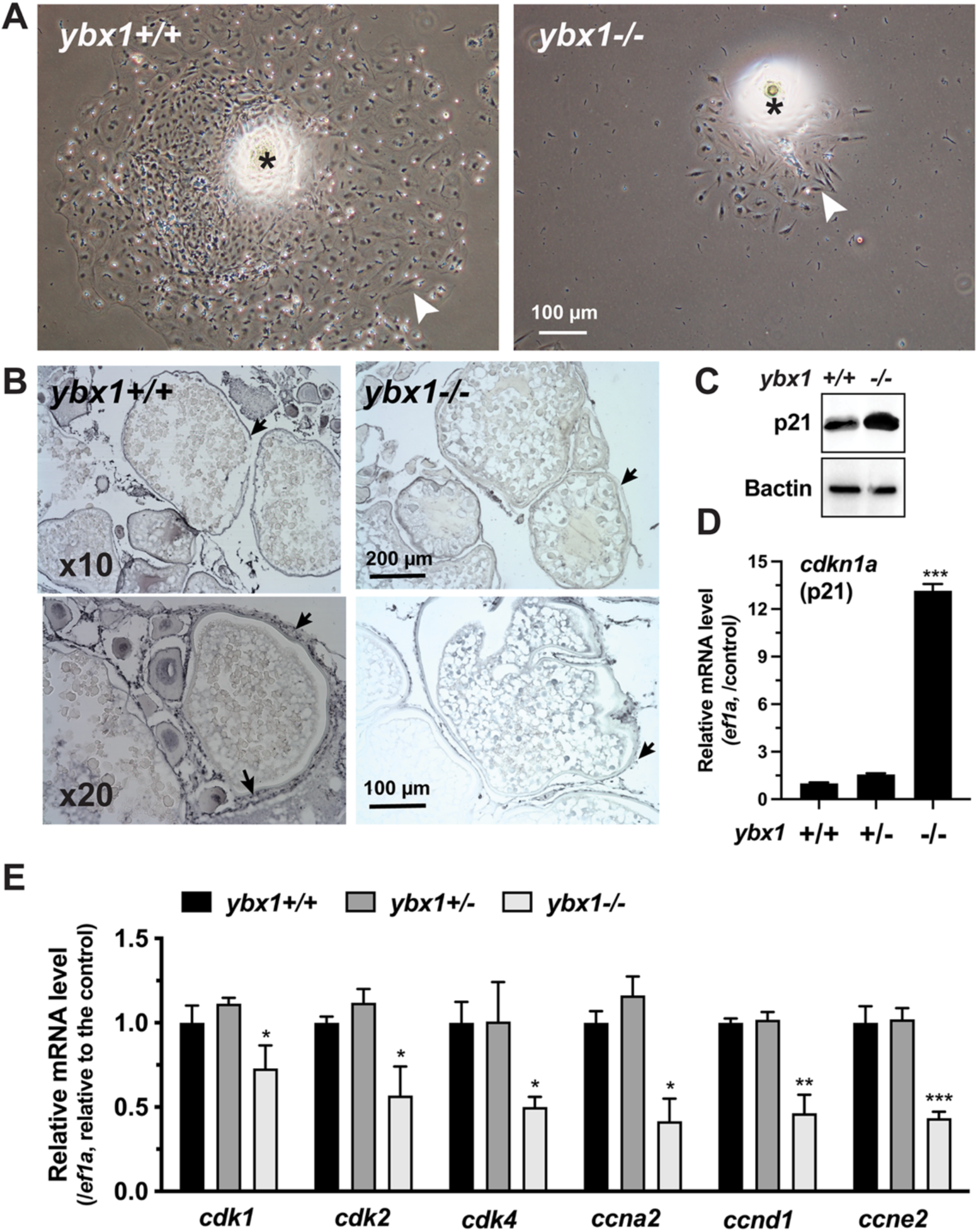
In vitro and in vivo assays on follicle cell proliferation. (A) Incubation of PV follicles from the control (*ybx1*+/+) and mutant (*ybx1*-/-) fish for 6 days. Asterisk, attached follicle; arrowhead, proliferated follicle cells. (B) BrdU incorporation assay. Arrow, follicle layer. (C) Western blot on p21 in PV follicles. (D) Real-time qPCR assay on *cdkn1a* (p21) expression in PV follicles. (E) Expression of cell cycle promoting factors in the PV follicles from the controls (*ybx1*+/+ and *ybx1*+/-) and mutant (*ybx1*-/-).

To further investigate the proliferation activity of follicle cells in vivo, we performed a BrdU incorporation assay. As shown in Fig. 5B, the control ovary (*ybx1*+/+) contained abundant BrdU-positive cells in both the follicle layer and inter-follicular space, indicating active somatic cell proliferation, including follicle cells. In sharp contrast, the *ybx1*-/- mutant ovary showed significantly fewer positive cells around the oocytes (Fig. 5B), indicating impaired proliferation of somatic cells in the *ybx1* mutant. This observation is consistent with the result obtained from the primary culture of follicle cells (Fig. 5A).

### Identification of p21 (cdkn1a) as a potential downstream target gene of Ybx1

To further confirm that the *ybx1*-/- follicles had impaired proliferation of follicle cells, which could be a mechanism for the blocked transition from PV to EV, or from previtellogenic to vitellogenic growth, we screened several genes that are potentially involved in regulating cell cycle or proliferation by Western blot, including Akt, MAPK, PCNA and p21 (data not shown). Among these proteins, p21 (*cdkn1a,* a potent cell cycle inhibitor) showed a remarkable increase in the PV follicles of *ybx1-/-* compared to the stage-matched control follicles (*ybx1+/+*) (Fig. 5C). Real-time qPCR also showed a marked increase of *cdkn1a* mRNA in the mutant follicles (*ybx1-/-*) compared with the stage-matched controls (*ybx1+/+* and *ybx1+/-*), further confirming the result with Western blot (Fig. 5D).

The increased expression of *cdkn1a* prompted us to analyze the expression of some other factors involved in cell cycle control. Interestingly, in contrast to *cdkn1a,* all the cell cycle-promoting examined showed decreased expression in the follicles of *ybx1-/-* mutant compared to the controls (*ybx1+/+* and *ybx1+/-*), including cyclin dependent kinases (*cdk1, cdk2* and *cdk4*) and cyclins (*ccna2*, *ccnd1* and *ccne2*) (Fig. 5E).

### *Mutagenesis of cdkn1a* (p21)

To investigate potential roles of p21 in Ybx1 actions, we used CRISPR/Cas9 method to disrupt the *cdkn1a* gene. A series of indel mutant lines were established and we chose the line with 14-bp deletion for phenotype analysis (Fig. S2A).

Survival analysis showed that *cdkn1a* deficiency resulted in severe mortality. Out of 3000 fish screened (> 2 mpf), we only obtained 8 homozygous mutant fish (*cdkn1a*-/-) (0.27% instead of the expected 25%). The ratio of the heterozygote (*cdkn1a*+/-) was also lower than the WT control (*cdkn1a*+/+), suggesting partial post-hatching lethality (Fig. S2B). To confirm the knockout of *cdkn1a* gene and provide further evidence for p21 involvement in folliculogenesis, we examined the expression of p21 protein in the control and mutant ovaries, which were genotyped by HRMA (Fig. S2C). Western blot analysis showed a complete lack of p21 in the mutant ovary (*cdkn1a*-/-). As expected, the level also decreased significantly in the heterozygote (*cdkn1a*+/-) compared with the control (*cdkn1a*+/+) (Fig. S2D).

Histological examination of *cdkn1a*-/- mutant ovaries at approximately 90 dpf demonstrated atypical ovarian architecture and disrupted follicle development. Relative to the control fish (*cdkn1a*+/+), the ovaries of the mutant exhibited a marked increase in inter-follicular somatic cells. Furthermore, oocytes within the mutant ovaries (*cdkn1a*-/-) were encased in an unusually thick chorion and lacked appropriately formed yolk granules (Fig. 6A).

**Fig 6.**
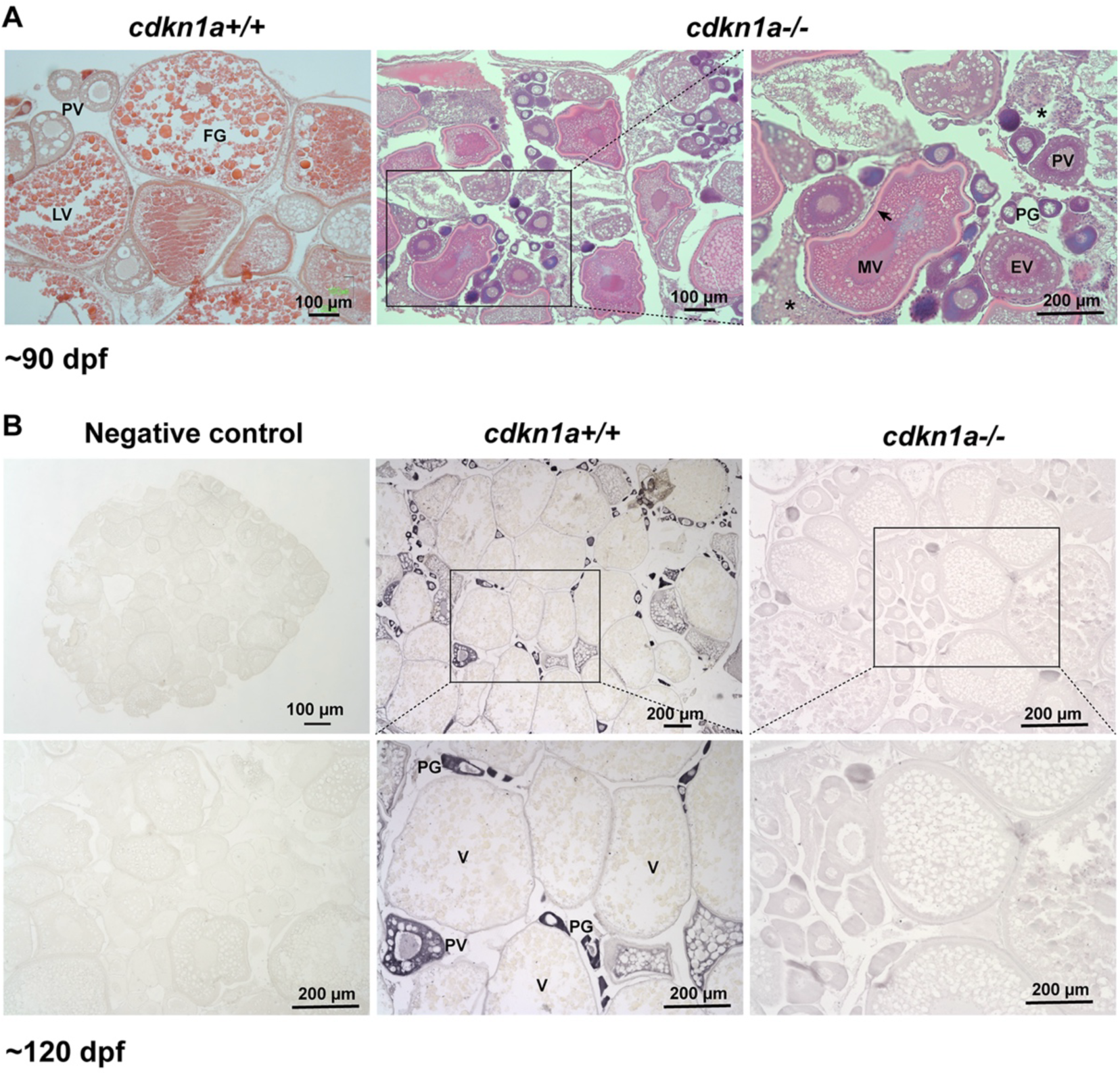
Spatiotemporal expression of *cdkn1a* (p21) in the ovary and the effect of *cdkn1a* mutation on follicle development. (A) Ovarian structure and follicle development in *cdkn1a* mutant around 90 dpf. (B) IHC staining for p21 protein in the control (*cdkn1a*+/+) and mutant (*cdkn1a*-/-) ovary around 120 dpf. PG, primary growth; PV, pre-vitellogenic; EV, early vitellogenic; MV, mid-vitellogenic; LV, late vitellogenic; FG, full-grown; V, vitellogenic follicles; arrow, thickened chorion; asterisk, inter- follicular somatic cells.

The successful knockout of *cdkn1a* gene was also confirmed by IHC staining, which detected no p21 protein in the ovary of *cdkn1a-/-* mutant. Interestingly, the spatiotemporal expression pattern of p21 protein in the control ovary (*cdkn1a+/+*) was surprisingly similar to that of Ybx1 as we reported recently (26). The p21 protein was predominantly expressed in the PG follicles (oocytes) with declining intensity at PV stages, and the signal vanished in vitellogenic follicles beyond PV stage (Fig. 6B).

### Precocious follicle activation in cdkn1a+/- females

Due to the severe lethality of the homozygous p21 mutant (*cdkn1a-/-*), we focused phenotype analysis on the heterozygote (*cdkn1a+/-*), which showed significantly higher survival rate despite partial lethality.

In control juvenile females (*cdkn1a+/+*), all follicles remained at PG stage when the body size (BL/BW) was below the threshold level for follicle activation or PG-PV transition (1.8 cm/100 mg) at 54 dpf, in agreement with our previous reports (34–38). However, follicle development was significantly advanced with premature follicle activation in the heterozygous mutant (*cdkn1a+/-*). PV follicles with cortical alveoli appeared much earlier in the mutant ovary when the body size was far below the threshold including 1.6 cm/71 mg in much younger fish (39 dpf), indicating advanced or premature follicle activation or puberty onset (Fig. 7A). Furthermore, the heterozygous mutant females also exhibited advanced vitellogenesis. At 50 dpf when the body sizes were within 2.2-2.3 cm (BL)/162-210 mg (BW), the ovaries contained full range of vitellogenic follicles from EV to FG stage with abundant yolk granules. In contrast, the size-matched control females (*cdkn1a+/+*; 2.1-2.3 cm/160-230 mg) contained PG ad PV follicles only without yolky follicles undergoing vitellogenic growth albeit at older age (65 dpf) (Fig. 7B). Quantification of follicles showed that the PV follicles first appeared in the heterozygous mutant ovary at BL 1.6 cm/BW 70-80 mg and the number increased significantly when the body size reached the threshold 1.8 cm/100 mg, when PV follicles just started to appear in the control ovary (Fig. 7C). FG follicles started to appear in the mutant ovary with body sizes around 2.1 cm/160- 180 mg whereas no vitellogenic follicles were observed in control females at 2.1-2.3 cm/160-230 mg (Fig. 7D). This agreed well with our recent study that 2.3 cm is the threshold BL for the development of vitellogenic follicles (EV-FG) (39).

**Fig 7.**
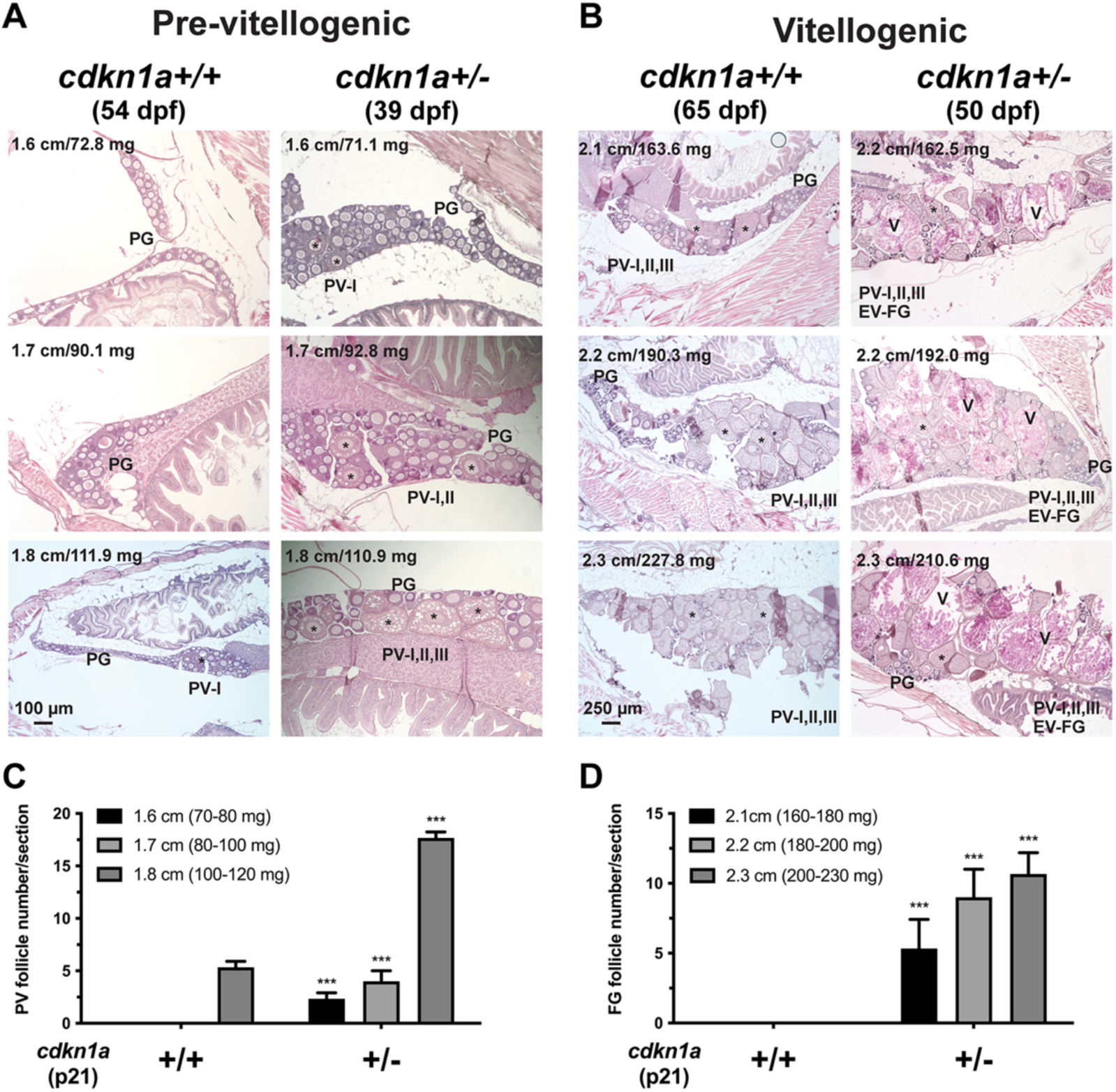
Effects of *cdkn1a* mutation on folliculogenesis in young fish. (A) Precocious follicle activation in *cdkn1a* heterozygous mutant (+/-) (39-54 dpf). PV follicles appeared earlier in the heterozygote than the size-matched WT control (+/+). (B) Advanced vitellogenic growth in the heterozygote (50-65 dpf). The mutant ovaries contained abundant vitellogenic follicles up to FG stage, whereas the size-matched control ovaries contained PV follicles only. PG, primary growth; PV-I, II, III, early, mid- and late PV follicles; EV-FG, early to full-grown vitellogenic follicles; V, vitellogenic follicles; asterisk, PV follicles. The body size (body length cm and weight mg) of each representative fish is shown in the photo. (C) Quantification of PV follicles in *cdkn1a* heterozygous mutant. (D) Quantification of FG follicles in the heterozygote. *** P < 0.001 compared with the size-matched control (n = 3).

### Premature ovarian failure in cdkn1a+/- mutant

Having demonstrated premature follicle activation in juvenile heterozygote (*cdkn1a+/-*) from 39 to 65 dpf, we went on to assess folliculogenesis in adult fish after sexual maturation from 3 to 8 months post-fertilization (mpf). The control fish displayed continuous active folliculogenesis throughout the study period at all time points. Folliculogenesis also occurred normally in the heterozygous fish up to 4 mpf; however, it ceased by 7 mpf with only PG and early PV follicles observable in the ovary. By 8 mpf, a subset of mutant fish (2 out of 11) exhibited a complete absence of PV follicles, retaining only early PG follicles in the ovary (Fig. 8A), which is reminiscent of premature ovarian failure in humans. The progressive deterioration of the ovary and its loss of function in the heterozygote were also reflected by the gradual decline of female fecundity from 2 to 8 mpf. The fish became almost infertile after 7 mpf (Fig. 8B).

**Fig 8.**
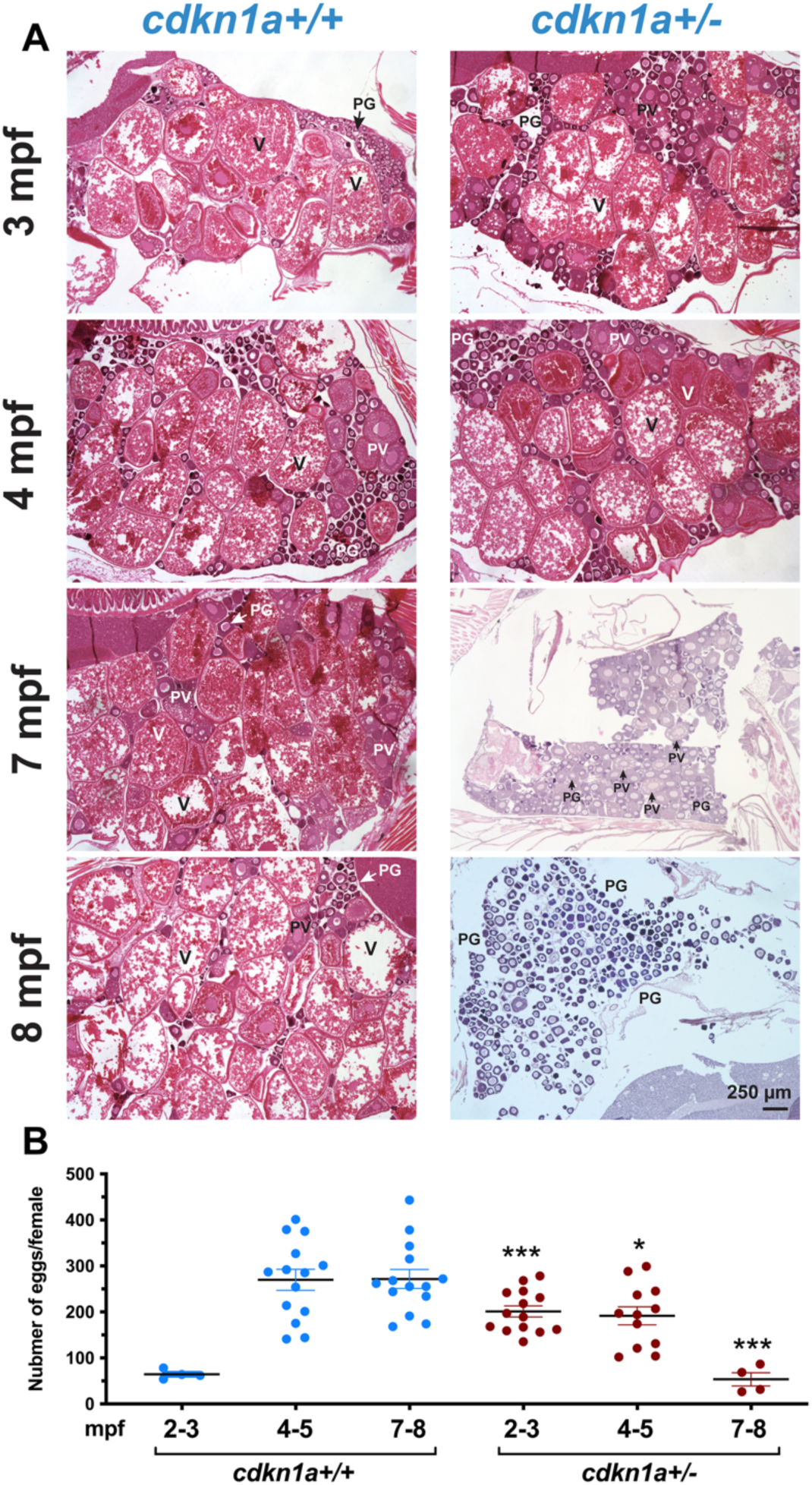
Effects of *cdkn1a* mutation on folliculogenesis in mature and elderly fish. (A) Ovarian histology of control (*cdkn1a*+/+) and heterozygous mutant (*cdkn1a*+/-) fish from 3 to 8 mpf. The heterozygous ovary showed normal folliculogenesis before 4 mpf; however, follicle development ceased from 7 mpf. PG, primary growth; PV, pre- vitellogenic; V, vitellogenic. (B) Fecundity (number of eggs spawned by each female) from 2 to 8 mpf. The data shown are mean ± SEM (n = 4-14). * P < 0.05, ***, P < 0.001 compared with stage-matched control.

### Transcriptome analysis for precocious follicle activation and premature ovarian failure in cdkn1a+/-

As described above, the heterozygous *cdkn1a+/-* mutants exhibited accelerated folliculogenesis with precocious follicle activation (PG-PV transition) in young fish, compared to the WT counterparts (*cdkn1a+/+*). To further investigate this phenomenon, we isolated PG follicles from *cdkn1a+/+* and *cdkn1a+/-* ovaries at 50 dpf for transcriptome analysis via RNA sequencing (RNA-seq). A substantial number of genes showed differential expression between the two groups (Fig. 9A). Interestingly, gene ontologies (GOs) pertaining to egg coat formation, oocyte development, gamete generation, and response to estradiol were significantly enriched and up-regulated in *cdkn1a+/-* follicles (Fig. 9B and C). Additionally, GOs associated with cell cycle regulation such as DNA replication initiation, pre-replicative complex assembly involved in nuclear cell cycle DNA replication, and double-strand break repair via break-induced replication were also up-regulated in *cdkn1a+/-* relative to the control. Conversely, the apoptotic signaling pathway was downregulated in *cdkn1a+/-* follicles (Fig. 9C). The KEGG pathway related to DNA replication was up-regulated in *cdkn1a+/-,* whereas the pathways of MAPK signaling, p53 signaling and cell cycle were down-regulated. The steroid biosynthesis pathway was enriched with both up- and down-regulated genes (Fig. 9D).

**Fig 9.**
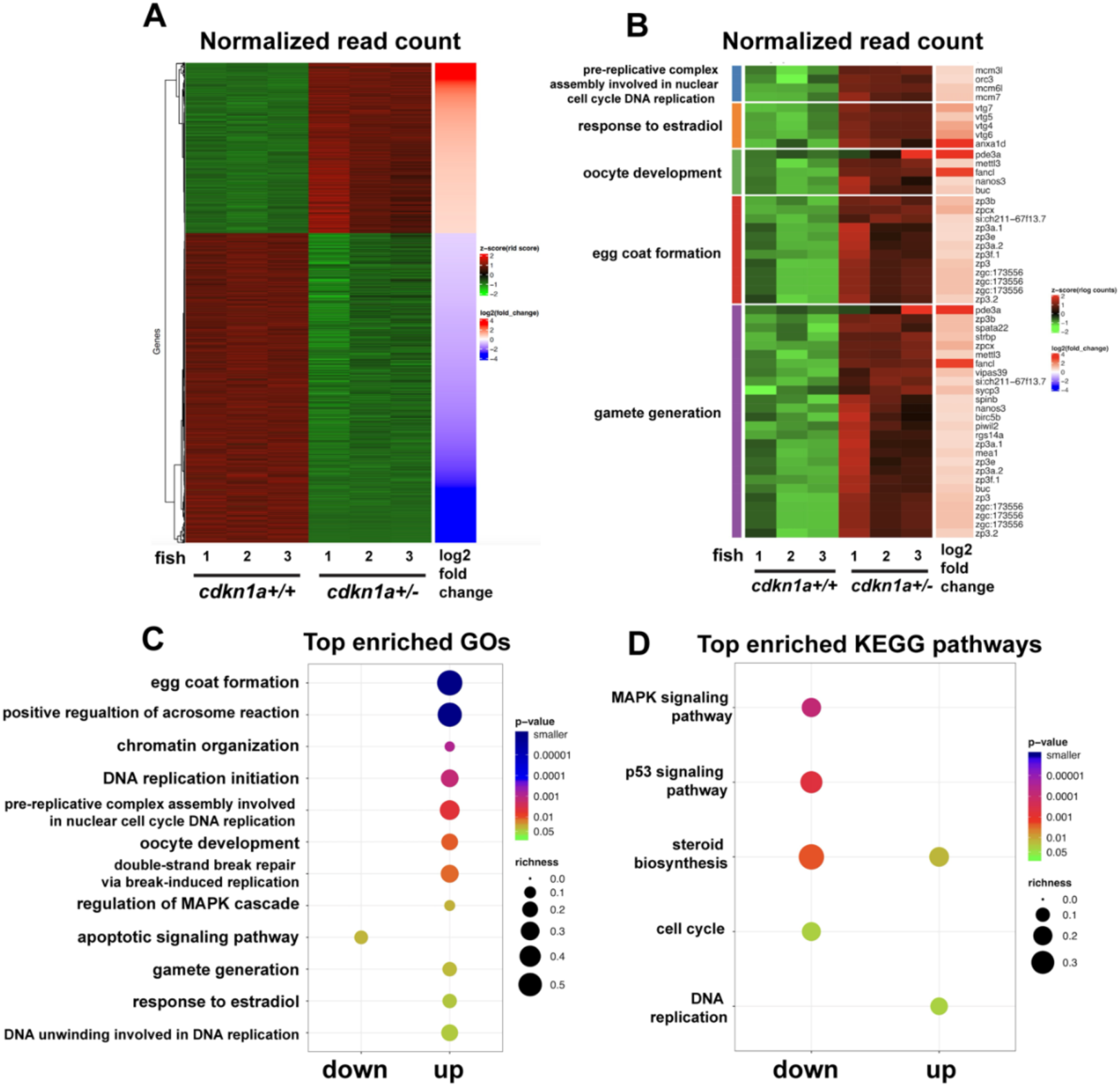
Transcriptome analysis on PG follicles from young control (*cdkn1a*+/+) and heterozygous fish (*cdkn1a*+/-) at 50 dpf. (A) Heatmap of DEGs between *cdkn1a*+/+ and *cdkn1a*+/-. (B) Heatmap of DEGs enriched in GOs associated with gamete development. (C) Top enriched GOs. (D) Top enriched KEGG pathways.

To provide clues to the mechanism underlying premature ovarian failure, we also performed transcriptome analysis on PG follicles from older *cdkn1a+/+* and *cdkn1a+/-* fish at 8 mpf when follicle degeneration occurred in *cdkn1a+/-* fish (Fig. 10A). Intriguingly, we found significantly diminished GOs associated with angiogenesis and cell growth in *cdkn1a+/-*, including vasculogenesis, lymphangiogenesis, sprouting angiogenesis, regeneration, and cell growth. However, the GOs of response to estradiol and cellular response to estrogen stimulus were significantly up-regulated in the mutant follicles (Fig. 10B and C). Additionally, KEGG pathways related to cell signaling such as Notch signaling pathway, Wnt signaling pathway, TGF-beta signaling pathway and MAPK signaling pathway were significantly enriched for down-regulated genes in *cdkn1a+/-* follicles, whereas the steroid biosynthesis pathway represents a mix of both up- and down-regulated genes (Fig.10D).

**Fig 10.**
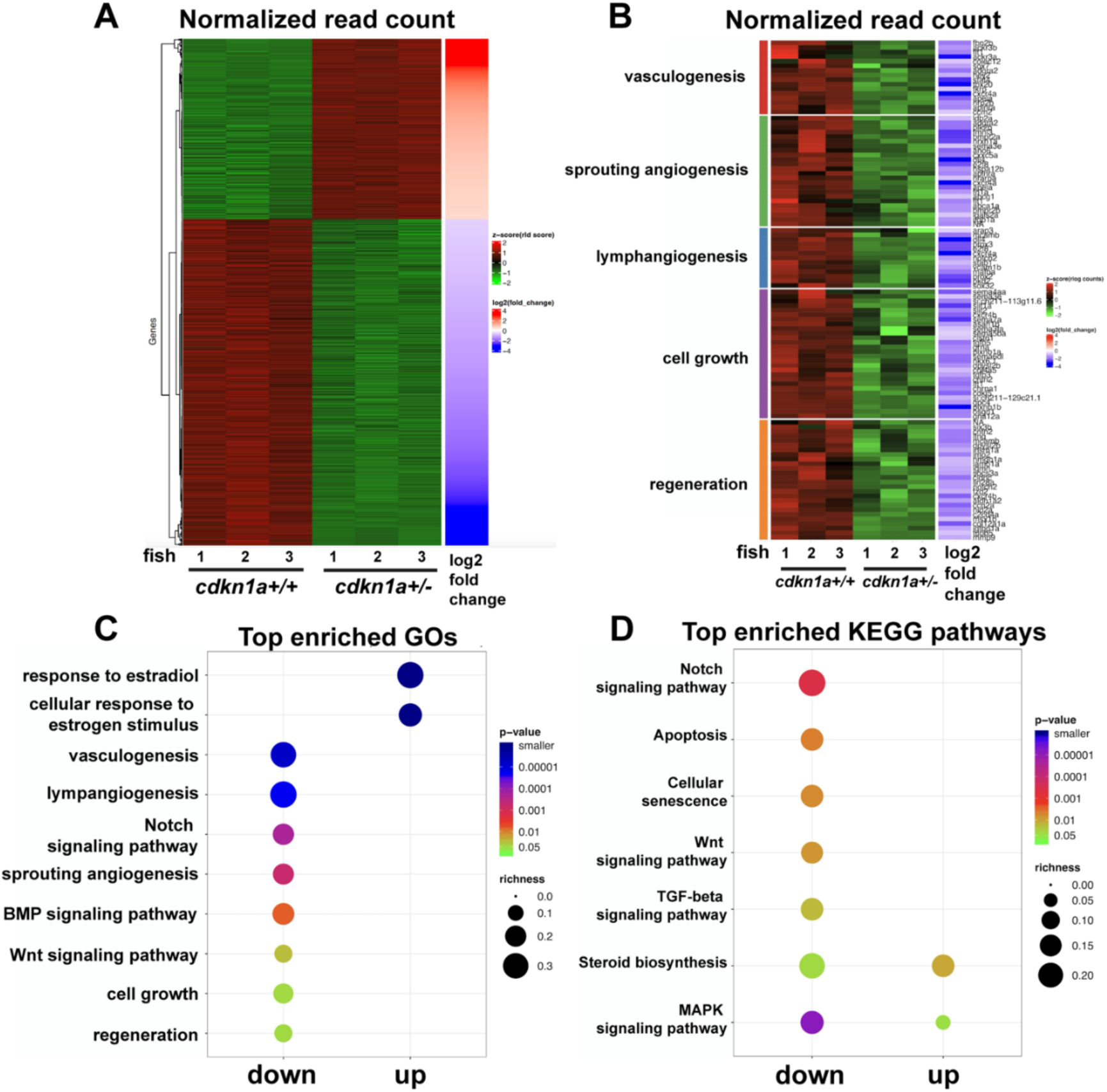
Transcriptome analysis on PG follicles from adult control (*cdkn1a*+/+) and heterozygous mutant (*cdkn1a*+/-) at 8 mpf. (A) Heatmap of DEGs between *cdkn1a*+/+ and *cdkn1a*+/-. (B) Heatmap of DEGs enriched in GOs associated with gamete development. (C) Top enriched GOs. (D) Top enriched KEGG pathways.

### Rescue of ovarian atrophy and follicle blockade in ybx1 mutant by simultaneous mutation of cdkn1a

The significant up-regulation of *cdkn1a* (p21) in the follicles of *ybx1* mutant (*ybx1*-/-) raised an interesting question about roles of p21 in Ybx1 action and the phenotypes of its mutation, in particular the arrest of follicle development at PV stage in the *ybx1*-/- mutant. To address this issue, we generated a double mutant of *ybx1* and *cdkn1a* (*ybx1-/-;cdkn1a-/-*). Due to the high mortality rates in both *ybx1-/-* (∼7% survival rate) and *cdkn1a-/-* (∼0.2% survival rate) single mutants, it was nearly impossible to obtain such double mutants for analysis. Fortunately, we obtained one female double mutant fish (∼4 mpf), which exhibited a surprising phenotype: rescue of follicle blockade in *ybx1-/-* females. Simultaneous loss of p21 could rescue the phenotypes of ovarian atrophy and follicle arrest in the *ybx1*-/- females as shown by the normal growth of the ovary in the double mutant. Histological analysis showed that follicle development, especially vitellogenic growth, resumed in the double mutant with abundant vitellogenic follicles present in the ovary, in contrast to the PV follicles only in the *ybx1-/-* single mutant ovary (Fig. 11).

**Fig 11.**
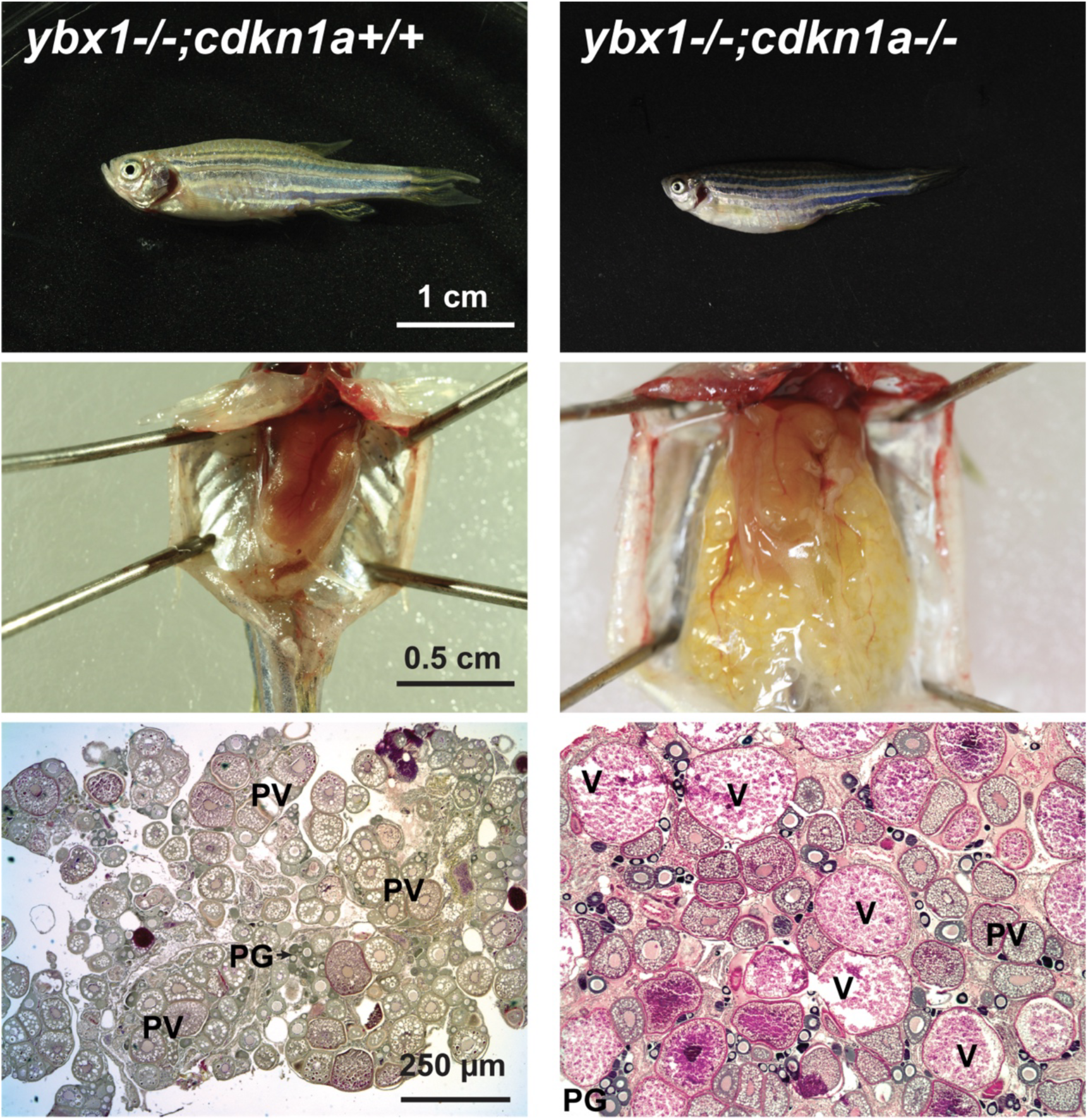
Rescue of *ybx1*-/- phenotype by simultaneous mutation of *cdkn1a*. One double mutant (*ybx1*-/-*;cdkn1a-/-*) was obtained around 4 mpf. Morphometric analysis showed normal growth of the ovary with normal folliculogenesis, in particular the appearance of vitellogenic follicles of different stages compared with the *ybx1*-/- single mutant whose follicles were blocked at PV stage. PG, primary growth; PV, pre-vitellogenic; V, vitellogenic.

### Rescue of impaired follicle cell proliferation in ybx1 mutant by mutation of cdkn1a

The surged expression of *cdkn1a* (p21) in *ybx1*-/- ovary and its primary function of inhibiting cell cycle led us to hypothesize that the mechanism by which *cdkn1a* mutation rescued *ybx1-/-* phenotypes of ovarian atrophy and follicle arrest might be related to cell proliferation in the follicle, particularly the somatic follicle cells whose proliferation is essential for folliculogenesis but impaired in *ybx1-/-* mutant.

To test this hypothesis and to substantiate the role of p21 in regulating follicle cell proliferation, we employed the same in vitro approach to explore the proliferative activity of follicle cells from various genotypes involving *ybx1* and *cdkn1a* (*ybx1+/+;cdkn1a+/+, ybx1+/+;cdkn1a+/-, ybx1-/-;cdkn1a+/+,* and *ybx1-/-;cdkn1a+/-*). Due to the high mortality rate, we used heterozygous *cdkn1a+/-* in different combinations. First, we isolated PV follicles from fish of these genotypes and incubated them for six days, during which time the follicles adhered to the dish and the follicle cells proliferated to form a circular ring of cell outgrowth around the oocyte. We then measured the diameters of these cell outgrowths as an indicator of follicle cell proliferation (Fig. 12A). Subsequently, we suspended the cultured follicle cells to remove the oocytes and sub-cultured them. The proliferative activity of the sub-cultured follicle cells was determined by MTT assay after three days of incubation (Fig. 12B). The results showed that both *ybx1+/+;cdkn1a+/+* (WT) and *ybx1+/+;cdkn1a+/-* follicle cells exhibited high proliferative activity, as evidenced by the formation of large growth rings. Interestingly, the *ybx1+/+;cdkn1a+/-* fish, which lost one *cdkn1a* allele, showed augmented proliferative activity. In contrast, the *ybx1* mutant (*ybx1-/-;cdkn1a+/+*) exhibited limited proliferative activity, with only a few follicle cells growing from the oocyte. However, partial loss of *cdkn1a* in *ybx1-/-;cdkn1a+/-* rescued the phenotype of *ybx1*-/-, as evidenced by the revived proliferation of follicle cells (Fig. 12A and C). These results were further confirmed by MTT assay on sub-cultured follicle cells (Fig. 12B and D). The presence and absence of Ybx1 proteins in cultured follicle cells were verified by Western blot, which showed a strong signal for Ybx1 in *ybx1+/+;cdkn1a+/+, ybx1+/+;cdkn1a+/-,* but not *ybx1-/-;cdkn1a+/+,* and *ybx1-/-;cdkn1a+/-* (Fig. 12E)

**Fig 12.**
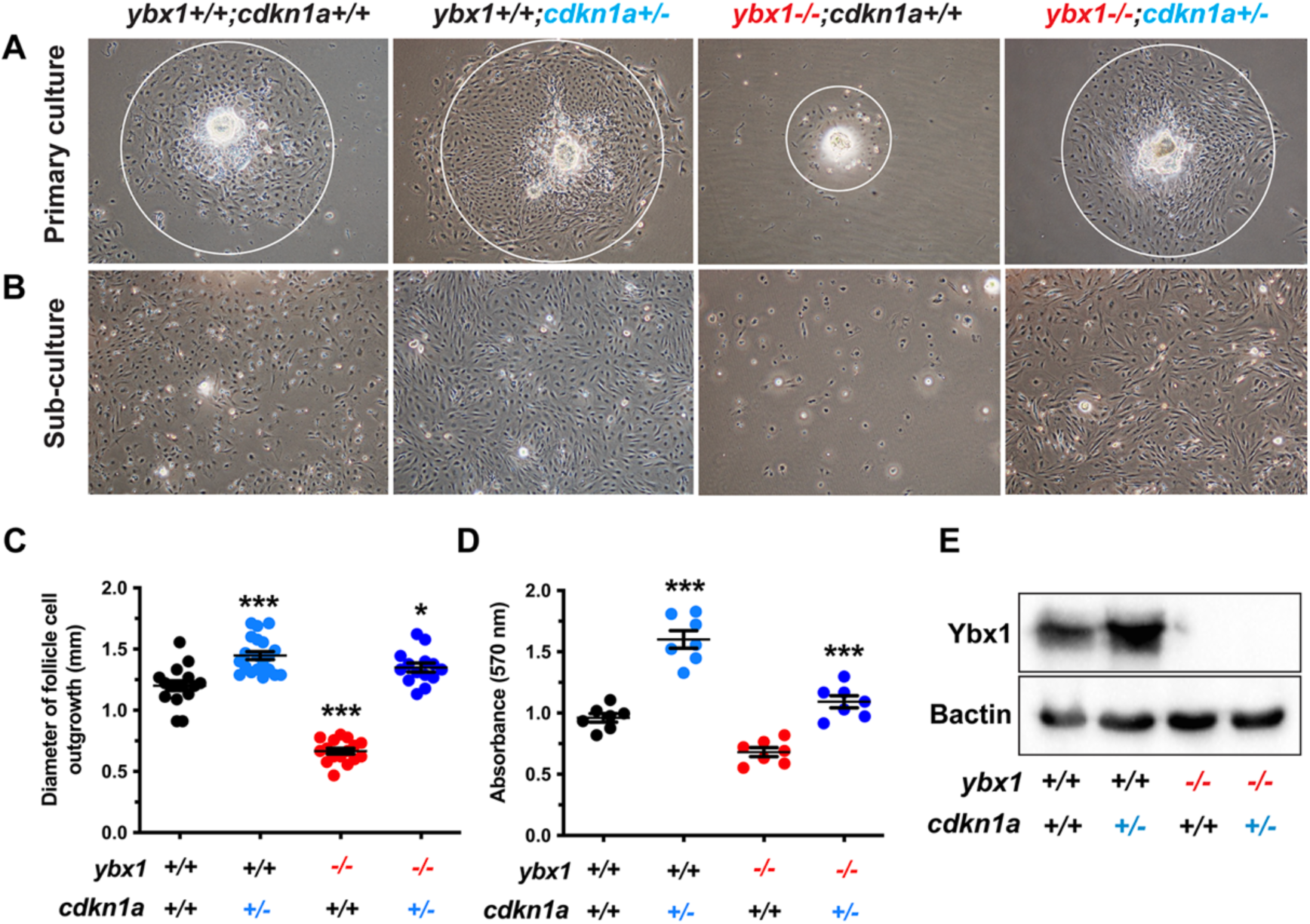
Rescue of impaired follicle cell proliferation by simultaneous mutation of *cdkn1a.* (A) Primary follicle culture for 6 days. The follicle cells proliferated actively from the attached follicle. (B) Sub-culture of the follicle cells after 6-day incubation. (C) Quantification of the follicle cell growth zone in different genotypic combinations (n = 14-20). (D) MTT assay on sub-cultured follicle cells from different genotypic combinations (n = 7). (E) Western blot on Ybx1 in cultured follicle cells of different genotypic combinations. The data shown are mean ± SEM. * P < 0.05, *** P < 0.001 compared with the control group.

## Discussion

Our recent proteomic analysis on zebrafish PG and PV follicles identified Ybx1 as the most abundant protein in the PG follicles. However, its level decreased significantly during the PG-PV transition when the follicles progressed to PV stage and continued to diminish afterwards to almost undetectable levels in subsequent stages of vitellogenic growth. This led us to hypothesize that Ybx1 could play a critical gate- keeping role in controlling the PG-PV transition or follicle activation (26). To test this hypothesis, we inactivated the *ybx1* gene in this study for functional analysis. Unfortunately, the mutant fish exhibited high post-hatching mortality rates, which constrained our analysis of reproductive performance in adults. Nevertheless, a small fraction of the mutant fish, roughly 7%, managed to survive to adulthood, which provided us an opportunity to assess the functional significance of Ybx1 in reproduction and further explore its regulatory mechanisms in folliculogenesis.

### Evidence for dual roles of Ybx1 in zebrafish

YB-1 belongs to the Y box-binding protein family in animals, which also includes other forms of molecules including YB-2 and YB-3 (1, 40). In contrast to YB- 1, which is expressed ubiquitously in various tissues (1), the expression of YB-2 is restricted to the gonads (41). In agreement with this, the loss of YB-1 (*Mys1*) in mice by gene knockout exhibited embryonic lethality, indicating functional importance of the protein in animal development (9). In contrast, the mutation of YB-2 (*Mys2*) did not affect the development and survival of the mutant mice; however, the mutants were infertile (18).

Unlike other vertebrates, zebrafish has only one form of YB proteins, namely YB-1 (Ybx1/ybx1). It is structurally homologous to mammalian YB-1, exhibiting ubiquitous mRNA expression. However, its protein is highly concentrated in the gonads, similar to mammalian YB-2 (26). This observation raises an interesting question: what would be the consequences of disrupting the *ybx1* gene in zebrafish? Would the resulting phenotypes of *ybx1*-/- fish mirror those of mammalian YB-1 or YB-2 mutants? Intriguingly, our current study revealed that the phenotypes of *ybx1*-/- mutant zebrafish bore similarities to both YB-1 and YB-2 mutants in mice. Firstly, *ybx1* gene inactivation resulted in high mortality, with only ∼7% of mutant individuals being able to survive to adulthood, a situation reminiscent of the YB-1 mutant in mice. Secondly, the survived female mutants exhibited sub-fertility, characterized by ovarian atrophy and markedly low fecundity, a trait similar to the YB-2 mutant in mice. These findings corroborate our proposed hypothesis that Ybx1 in zebrafish may serve dual functional roles, analogous to both YB-1 and YB-2 (26).

### Ybx1 is not required for the transition from primary to secondary growth

As we reported recently and observed in the present study, there was an abundance of Ybx1 protein in PG follicles, which decreased significantly when the follicles entered the PV stage (PG-PV transition). This led us to hypothesize that Ybx1 might play a pivotal role in controlling this transition or follicle activation (26). However, the phenotypic analysis of the *ybx1*-/- mutant in the current study does not support this hypothesis, demonstrating that Ybx1 is not essential for follicle activation or the PG-PV transition from the primary to secondary growth phase. Surprisingly, PV follicles could develop normally in *ybx1* mutant with formation of cortical alveoli in the oocytes in the absence of Ybx1. This finding implies that the dramatic decrease of Ybx1 protein at the PV stage, relative to its prevalence in PG follicles, is not vital for the development of PV follicles per se. Instead, it may play a role in subsequent stages of follicle development.

### Ybx1 is required for the transition from previtellogenic to vitellogenic growth

After determining that Ybx1 does not contribute to follicle activation or the PG-PV transition, we shifted our attention to the subsequent stages of the secondary growth phase, specifically vitellogenic growth. Interestingly, the absence of Ybx1 caused follicles to arrest at the PV stage, thereby preventing their progression to vitellogenic growth, which commences at the early vitellogenic (EV) stage. This disruption resulted in significantly smaller ovaries (atrophy) in the mutant compared to controls.

PV follicles are characterized with the formation of cortical alveoli without yolk granules, whereas EV follicles begin accumulating yolk granules around the germinal vesicle (GV) in the oocyte, marking the onset of vitellogenic growth (35–37, 42). Unexpectedly, despite the majority of follicles being stuck at the PV-EV transition, a small number managed to bypass this blockage at later ages and initiate vitellogenic growth. These follicles could ultimately undergo final oocyte maturation and ovulation. Consequently, the *ybx1*-/- mutant females could breed with WT males to produce offspring, but with a significantly reduced fecundity.

Intriguingly, the phenotype of follicle arrest at PV stage in the *ybx1*-/- mutant mirrors that of the bone morphogenetic protein 15 (*bmp15*-/-) mutant (35, 43). Given that both *bmp15* and *ybx1* are predominantly expressed in the oocyte (26, 35), the phenotypic similarity of their mutants raises an interesting question about their potential functional or regulatory relation. In a recent study, we demonstrated that Ybx1 was a major component of the messenger ribonucleoprotein particles (mRNPs) in the zebrafish follicles, and it was associated with mRNAs of multiple growth factors, particularly those expressed in the oocyte like growth differentiation factor 9 (*gdf9*) and members of the BMP family (26). Whether Ybx1 also binds to Bmp15 mRNA and thus phenocopies each other when inactivated remains an interesting question that warrants further exploration in future studies.

### Impaired follicle cell proliferation in ybx1-/- mutant

Follicle development in fish is propelled by both oocyte growth and proliferation of the somatic follicle cells encircling the oocyte (44, 45). To explore the mechanisms that underpin the blockade of follicle growth in *ybx1*-/- fish, we evaluated the proliferative potential or capability of the mutant follicle cells by using a novel follicle cell culture system we previously developed (32). This method has been widely used to investigate the regulation of various genes expressed in the follicle cells (29, 46–50). Our results revealed that the follicle cells of the *ybx1*-/- PV follicles have lost their ability to proliferate, both in vivo and in vitro, in sharp contrast to those of the WT fish. This could be one of the underpinning mechanisms for follicle arrest in the *ybx1*-/- mutant.

The decline in follicle cell proliferative ability in *ybx1*-/- mutant follicles agrees well with the reports that YB-1 promotes cell proliferation in a variety of cell types, especially cancer cells (51). In human melanoma and breast cancer stem cells, disruption of YB-1 gene with CRISPR/Cas9 inhibited cell proliferation, leading to cell cycle arrest (52), similar to our observation of impaired follicle cell proliferation in *ybx1*-/- mutant. Evidence has accumulated that YB-1 stimulates cell cycle progression by activating multiple pro-proliferation genes including cyclins and epidermal growth factor receptor (EGFR) (51). Consistent with this, mutation of YB-1 or knocking down its expression suppressed expression of EGFR and therefore cell proliferation in breast tumor cells (53).

### Role of p21 in zebrafish folliculogenesis

The arrest of follicles at PV stage in the *ybx1-/-* mutant coincided with a loss of proliferative ability of the follicle cells. This prompted us to turn our attention to the regulation of cell cycle and proliferation. Through Western blot screening of potential proteins associated with cell proliferation, we observed a remarkable increase in the protein level of p21 (*cdkn1a*) in *ybx1-/-* mutant follicles, which was also confirmed by real-time qPCR. Our IHC staining showed clearly that p21 was present in the zebrafish ovary, with an expression pattern similar to that of Ybx1. The p21 protein level was peaked in the PG follicles, but significantly decreased during PG-PV transition and continued to diminish thereafter. This suggests that p21, like Ybx1, may also play a role in controlling folliculogenesis, similar to Ybx1.

p21, also known as Cip1, Waf1, Sdi1, Mda6 and Cap20, is a well-established cell cycle inhibitor that halts cells at G1 or S phase by inhibiting a broad spectrum of cyclin-dependent kinases (CDKs) that drive the cell cycle (54). The expression of p21 is subject to regulation by various exogenous and endogenous factors that regulate or interfere with cell cycle (55). Overexpression of p21 induces cell cycle arrest in numerous cell lines (56), while suppression or depletion of p21 expression promotes cell cycle progression and DNA synthesis (57). Given these known functions of p21 and its elevated expression in *ybx1*-/- follicles, we hypothesized that the increased expression of p21 might be accountable for the ceased proliferation of the follicle cells in the *ybx1-/-* mutant.

To further substantiate the involvement of p21 in Ybx1-controlled follicle cell proliferation, we disrupted the p21 gene *cdkn1a* with CRISPR/Cas9 in zebrafish. Complete loss of p21 in the homozygous mutant (*cdkn1a-/-*) resulted in severe post-hatching mortality, a stark contrast to the p21 null mutant in mice, which exhibited normal development and survival rates. The primary abnormality observed in the p21-/- mutant mice was a deficient ability of the mutant embryonic fibroblasts to arrest at G1 stage of the cell cycle (58).

Interestingly, although the zebrafish heterozygote (*cdkn1a+/-*) could survive well into the adulthood, its follicles showed precocious activation and maturation in young zebrafish. This phenotype contrasted with that of the *ybx1-/-* mutant, whose follicles arrested at the PV stage. However, the *cdkn1a+/-* ovaries began to exhibit signs of premature ovarian failure after 4 mpf, and the follicle development ceased completely after about 7-8 mpf. These phenotypes strongly suggest that p21 may serve as a vital regulator controlling the initiation and progression of folliculogenesis.

To understand the potential mechanisms underlying the precocious follicle activation (puberty onset) in young fish and premature ovarian failure in adults, we performed transcriptome analyses on PG follicles from *cdkn1a+/+* and *cdkn1a+/-* at 50 dpf and 8 mpf (240 dpf) respectively. Interestingly, a large number of DEGs were up-regulated in *cdkn1a+/-* fish compared to *cdkn1a+/+* at 50 dpf, and many were enriched in GOs associated with cell cycle and gametogenesis (folliculogenesis). These genes are supposed to promote and accelerate follicle development, and their up- regulation may be underpinning the precocious follicle activation in young zebrafish. In contrast, many DEGs down-regulated in *cdkn1a+/-* fish at 8 mpf were enriched in GOs linked with cell growth and angiogenesis. This aligns well with our previous report that the biological processes or pathways associated with angiogenesis were all enriched by up-regulated DECs during the PG-PV transition or follicle activation (30).

### Role of p21 in Ybx1-controlled follicle cell proliferation

The complete loss of p21 (*cdkn1a-/-*) caused post-hatching lethality. Unexpectedly, experiments with the few surviving mutant fish revealed that the loss of p21 (*cdkn1a-/-*) could rescue the phenotype of *ybx1-/-* mutant, with follicles overcoming the PV blockade in the double mutant (*ybx1-*/-;*cdkn1a-/-*). Interestingly, the heterozygote (*cdkn1a+/-*) could also ameliorate the defects of folliculogenesis in the *ybx1-/-* mutant, particularly the impaired proliferative ability of the follicle cells. These have led us to hypothesize that Ybx1 plays a critical role in controlling early follicle development, especially the transition from previtellogenic to vitellogenic growth, and it achieves this, in part, by suppressing *cdkn1a* expression. This accelerates the proliferation of the somatic follicle cells, which is essential to support rapid follicle growth.

As an RNA/DNA binding protein, YB-1 serves dual roles: acting as a transcription factor to regulate gene expression and functioning as an RNA-binding protein to stabilize mRNAs while inhibiting their translation (1). The remarkable upsurge of p21 (*cdkn1a*) at both mRNA and protein levels in *ybx1*-/- follicles implies that Ybx1 may regulate its expression at both transcription and translation levels. It has been well established that p21 expression is subject to regulation by a variety of mRNA binding proteins (RBPs), which is a critical rate-limiting mechanism that determines the level of p21 in cells (55). As we reported recently, Ybx1 also functions as a key component of the messenger ribonucleoprotein particles (mRNPs) in zebrafish, and it binds mRNAs of a wide range of molecules in zebrafish follicles (26). It is possible that Ybx1 also binds *cdkn1a* mRNA and prevents its translation in zebrafish follicles.

It is worth noting that both Ybx1 and p21 are co-localized in the early follicles (PG and PV), primarily in the oocyte, and they also displayed similar temporal patterns of expression during folliculogenesis. Whether a regulatory interaction between Ybx1 and p21 occurs in the oocyte, the follicle cells, or both, remains to be determined. Our data on proliferation of cultured follicle cells, which are devoid of oocytes, suggest that the regulation occurs in the follicle cells. This also aligns well with our IHC staining results, which showed presence of Ybx1 in both oocytes and follicle layers.

The involvement of p21 in YB-1-regulated cell proliferation has been suggested in various studies, including those in mice. Disruption of the YB-1 gene (*Msy1*) in mice resulted in severe growth retardation during late embryogenesis, which was accompanied by increased expression of cyclin-dependent kinase inhibitors p16 and p21. These findings suggest an essential role for YB-1 in regulating cell proliferation, which is critical for normal embryogenesis (8). Consistent with this, embryonic fibroblasts derived from YB-1 mutants showed reduced growth and cell density in culture, indicating again that YB-1 is required for cell proliferation (9). Moreover, YB-1 may also act as an extracellular mitogen that activates cell migration and proliferation in primary cell culture (59). These findings agree well with our observations in the present study on zebrafish follicle development. A recent study in zebrafish showed that the nuclear translocation of Ybx1 was regulated by the circadian clock, and that Ybx1 interacted directly with the promoter of cyclin A2 to downregulate its expression (60). Interestingly, the expression of p21 (*cdkn1a*) in zebrafish is also subject to regulation by the circadian clock, showing a robust circadian oscillation (61). These findings support a role for Ybx1 in controlling cell cycle in zebrafish and the potential participation of p21 in the regulation. Our study found that the loss of Ybx1 led to a significant increase in the expression of cell cycle inhibitor p21 (*cdkn1a*) and a decrease in the expression of cyclins (*ccna2, ccnd1*, and *ccne2*) and cyclin-dependent kinases (*cdk1, cdk2*, and *cdk4*) in PV follicles. These results support the notion that YB-1 plays a key role in controlling cell proliferation by suppressing cell cycle inhibitors while enhancing cell cycle promoters, as previously reported (62).

## Conclusion and perspectives

In this study, we utilized genome editing technology to disrupt the *ybx1* gene in zebrafish, with the purpose of investigating its potential role in reproduction, particularly folliculogenesis in females. Contrary to our initial hypothesis, which proposed a critical role for Ybx1 in follicle activation or PG-PV transition, the results from the *ybx1*-/- mutant indicated otherwise. Despite the precipitous drop in Ybx1 level at the PG-PV transition, the PV follicles could form normally with cortical alveoli in the oocytes. Nonetheless, further analysis revealed that follicle development was arrested during the transition from previtellogenic to vitellogenic growth, also known as the PV-EV transition. We attributed this developmental arrest, at least partially, to an impairment in follicle cell proliferation. Further investigation revealed that the diminished proliferative activity of the follicle cells in *ybx1* mutant follicles was a result of an upsurge in the expression of the cell cycle inhibitor p21 (*cdkn1a*). By disrupting the *cdkn1a* gene, we were able to rescue the phenotypes of the *ybx1* mutant in terms of both folliculogenesis and follicle cell proliferation. The findings of this study provide robust evidence for the gatekeeping role of Ybx1 in the early stages of follicle development and its action mechanisms.

## Acknowledgement

We thank Ms. Phoenix Un Ian LEI for the maintenance and management of the zebrafish facility and the Genomics and Bioinformatics Core of the Faculty of Health Sciences for technical support. This study was supported by grants from the University of Macau (MYRG2020-00192-FHS, MYRG2022-00219-FHS, CPG2022-00028-FHS, and CPG2023-00029-FHS) and The Macau Fund for Development of Science and Technology (FDCT0132/2019/A3 and NSFC-FDCT0086/2022/AFJ) to WG.

---

**Fig S1.**
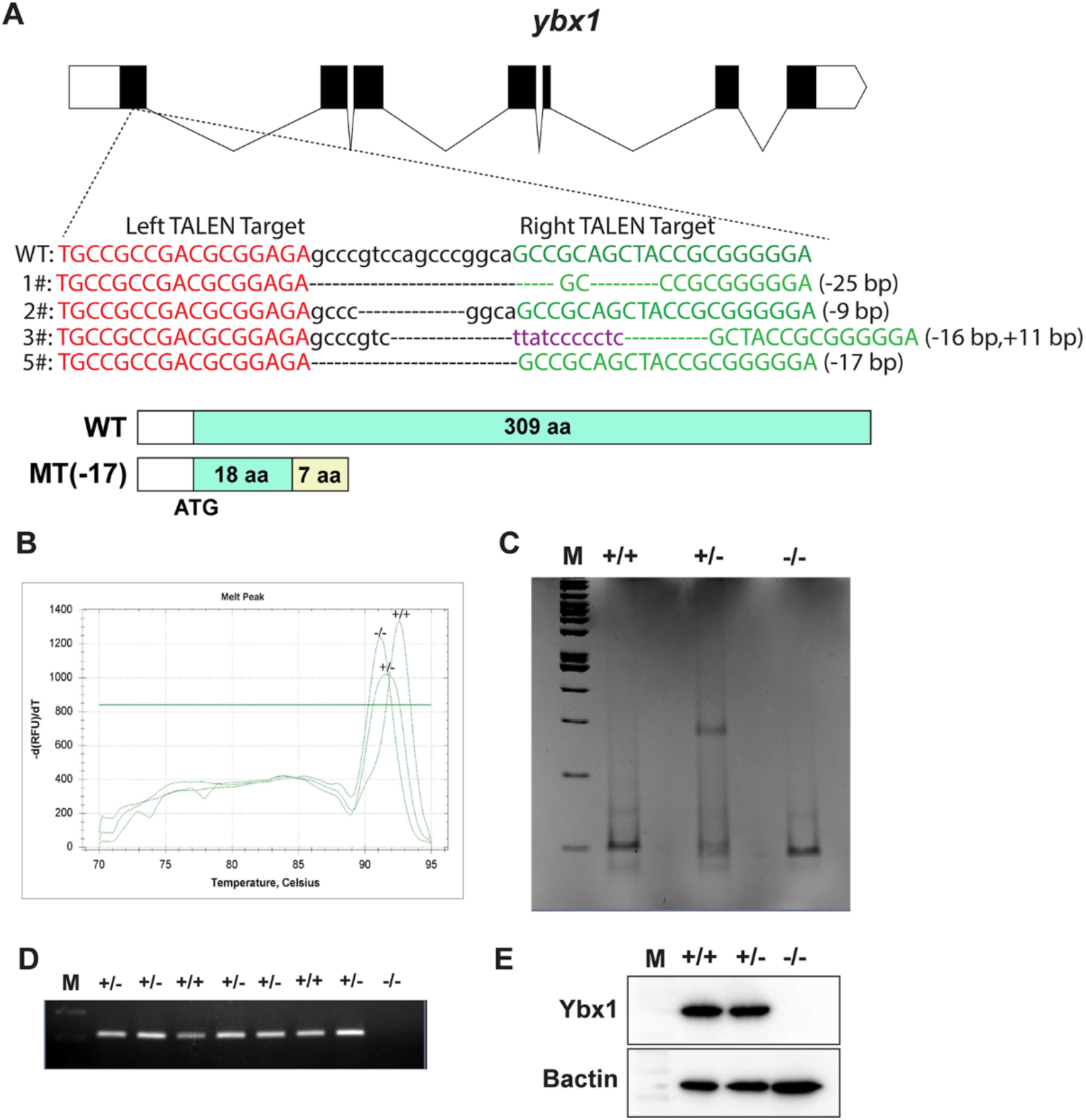
Targeted disruption of *ybx1* gene by TALEN. (A) Genomic structure of zebrafish *ybx1* gene. Several mutant lines were established and the line with 17-bp deletion (-17) was chosen for phenotype analysis. The mutation resulted in a premature termination in protein translation, producing a truncated protein with 18 amino acids. (B) HRMA assay showing different melt curves for three genotypes (+/+, +/-, and -/-). (C) HMA assay showing different banding patterns for three genotypes. (D) PCR detection of WT *ybx1* gene sequence in different genotypes with a mutant-specific primer. (E) Western blot on Ybx1 in ovaries of different genotypes.

**Fig S2.**
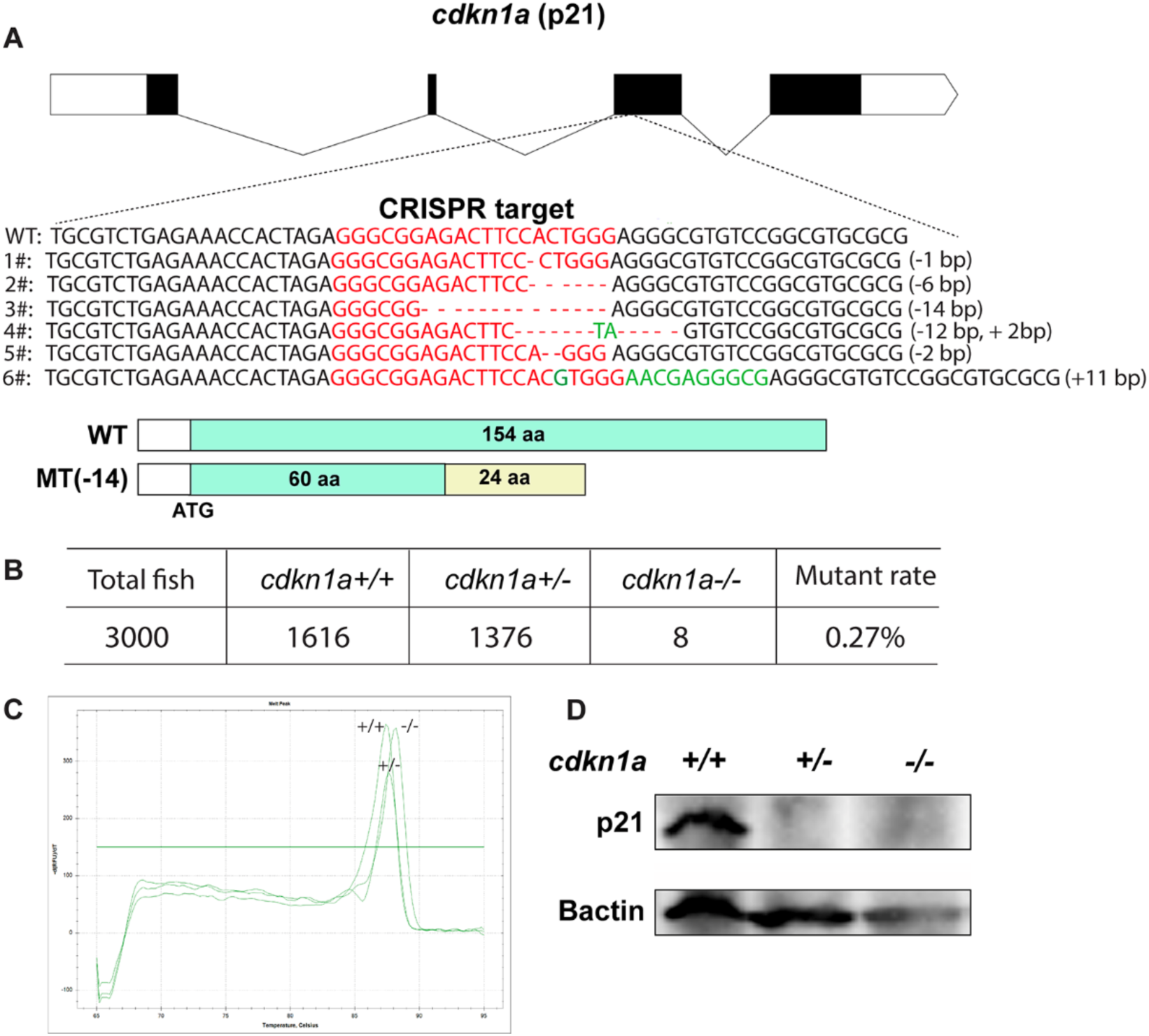
Targeted disruption of *cdkn1a* gene by CRISPR/Cas9. (A) Genomic structure of zebrafish *cdkn1a* gene. Several mutant lines were established and the line with 14- bp deletion (-14) was chosen for phenotype analysis. The mutation resulted in a premature termination in protein translation, producing a truncated protein with 60 amino acids. (B) Data on the survival rates of heterozygous and homozygous mutants of *cdkn1a*. (C) HRMA assay showing different melt curves for three genotypes (+/+, +/-, and -/-). (D) Western blot on p21 in ovaries of different genotypes.

**Supplementary Table S1:**
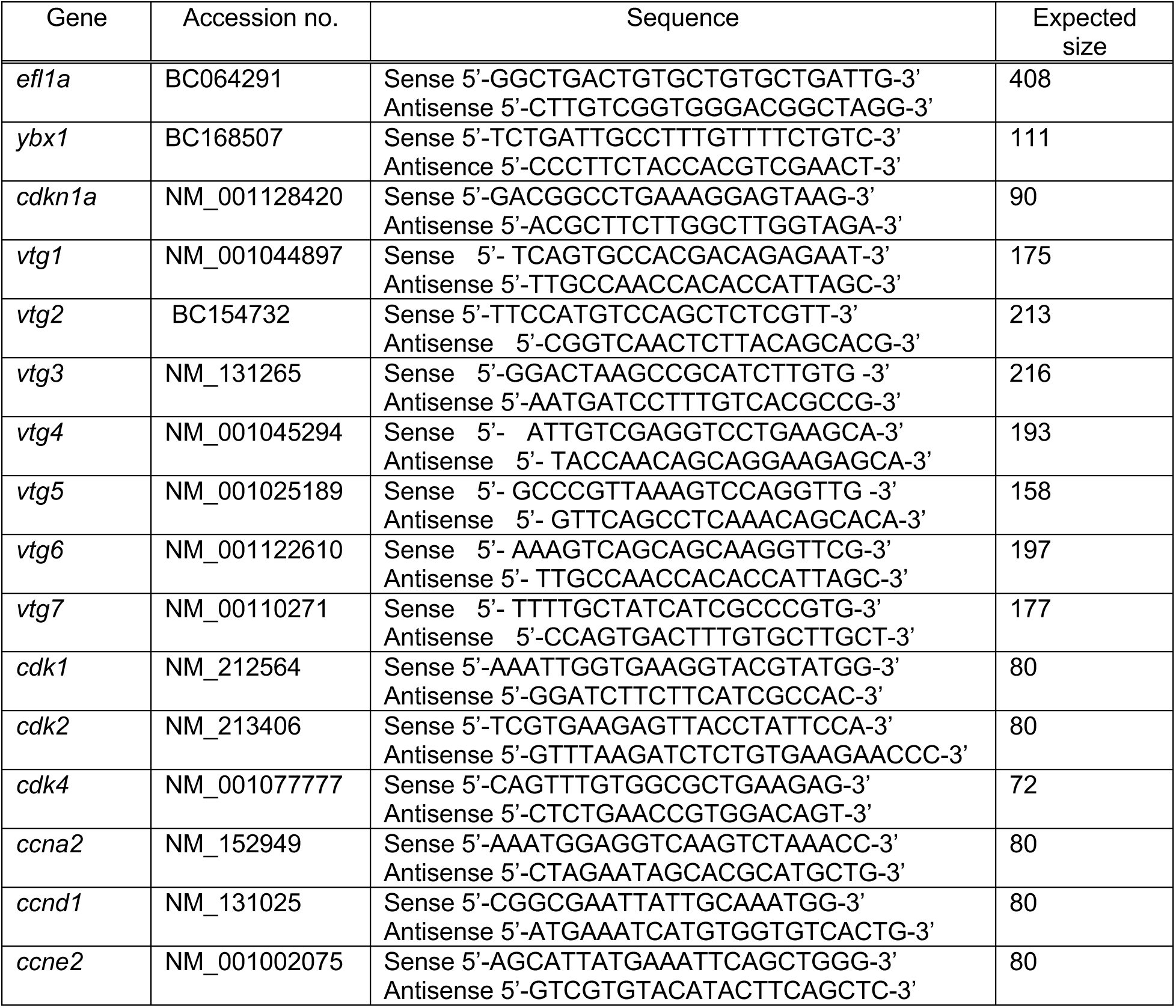

## References

1. Eliseeva IA, Kim ER, Guryanov SG, Ovchinnikov LP, Lyabin DN. Y-box-binding protein 1 (YB-1) and its functions. Biochemistry (Mosc) 2011; 76:1402–1433

2. Lyabin DN, Eliseeva IA, Ovchinnikov LP. YB-1 protein: functions and regulation. Wiley Interdiscip Rev RNA 2014; 5:95–110

3. Tafuri SR, Wolffe AP. Xenopus Y-box transcription factors: molecular cloning, functional analysis and developmental regulation. Proc Natl Acad Sci USA 1990; 87:9028–9032

4. Izumi H, Imamura T, Nagatani G, Ise T, Murakami T, Uramoto H, Torigoe T, Ishiguchi H, Yoshida Y, Nomoto M, Okamoto T, Uchiumi T, Kuwano M, Funa K, Kohno K. Y box-binding protein-1 binds preferentially to single-stranded nucleic acids and exhibits 3’-->5’ exonuclease activity. Nucleic Acids Res 2001; 29:1200–1207

5. Kohno K, Izumi H, Uchiumi T, Ashizuka M, Kuwano M. The pleiotropic functions of the Y-box-binding protein, YB-1. Bioessays 2003; 25:691–698

6. Yadav BS, Singh S, Shaw AK, Mani A. Structure prediction and docking-based molecular insights of human YB-1 and nucleic acid interaction. J Biomol Struct Dyn 2016; 34:2561–2580

7. Rauen T, Raffetseder U, Frye BC, Djudjaj S, Muhlenberg PJ, Eitner F, Lendahl U, Bernhagen J, Dooley S, Mertens PR. YB-1 acts as a ligand for Notch-3 receptors and modulates receptor activation. J Biol Chem 2009; 284:26928–26940

8. Lu ZH, Books JT, Ley TJ. YB-1 is important for late-stage embryonic development, optimal cellular stress responses, and the prevention of premature senescence. Mol Cell Biol 2005; 25:4625–4637

9. Uchiumi T, Fotovati A, Sasaguri T, Shibahara K, Shimada T, Fukuda T, Nakamura T, Izumi H, Tsuzuki T, Kuwano M, Kohno K. YB-1 is important for an early stage embryonic development: neural tube formation and cell proliferation. J Biol Chem 2006; 281:40440–40449

10. Kumari P, Gilligan PC, Lim S, Tran LD, Winkler S, Philp R, Sampath K. An essential role for maternal control of Nodal signaling. Elife 2013; 2:e00683. doi:00610.07554/eLife.00683

11. Sun J, Yan L, Shen W, Meng A. Maternal Ybx1 safeguards zebrafish oocyte maturation and maternal-to-zygotic transition by repressing global translation. Development 2018; 145:dev166587. doi:166510.161242/dev.166587

12. Arnold A, Rahman MM, Lee MC, Muehlhaeusser S, Katic I, Gaidatzis D, Hess D, Scheckel C, Wright JE, Stetak A, Boag PR, Ciosk R. Functional characterization of C. elegans Y-box-binding proteins reveals tissue-specific functions and a critical role in the formation of polysomes. Nucleic Acids Res 2014; 42:13353–13369

13. Mansfield JH, Wilhelm JE, Hazelrigg T. Ypsilon Schachtel, a Drosophila Y-box protein, acts antagonistically to Orb in the oskar mRNA localization and translation pathway. Development 2002; 129:197–209

14. Miwa A, Higuchi T, Kobayashi S. Expression and polysome association of YB- 1 in various tissues at different stages in the lifespan of mice. Biochim Biophys Acta 2006; 1760:1675–1681

15. Yang J, Medvedev S, Yu J, Schultz RM, Hecht NB. Deletion of the DNA/RNA-binding protein MSY2 leads to post-meiotic arrest. Mol Cell Endocrinol 2006; 250:20–24

16. Davies HG, Giorgini F, Fajardo MA, Braun RE. A sequence-specific RNA binding complex expressed in murine germ cells contains MSY2 and MSY4. Dev Biol 2000; 221:87–100

17. Gu W, Tekur S, Reinbold R, Eppig JJ, Choi YC, Zheng JZ, Murray MT, Hecht NB. Mammalian male and female germ cells express a germ cell-specific Y-Box protein, MSY2. Biol Reprod 1998; 59:1266–1274

18. Yang J, Medvedev S, Yu J, Tang LC, Agno JE, Matzuk MM, Schultz RM, Hecht NB. Absence of the DNA-/RNA-binding protein MSY2 results in male and female infertility. Proc Natl Acad Sci USA 2005; 102:5755–5760

19. Yu J, Hecht NB, Schultz RM. Expression of MSY2 in mouse oocytes and preimplantation embryos. Biol Reprod 2001; 65:1260–1270

20. Yang J, Morales CR, Medvedev S, Schultz RM, Hecht NB. In the absence of the mouse DNA/RNA-binding protein MSY2, messenger RNA instability leads to spermatogenic arrest. Biol Reprod 2007; 76:48–54

21. Medvedev S, Yang J, Hecht NB, Schultz RM. CDC2A (CDK1)-mediated phosphorylation of MSY2 triggers maternal mRNA degradation during mouse oocyte maturation. Dev Biol 2008; 321:205–215

22. Medvedev S, Pan H, Schultz RM. Absence of MSY2 in mouse oocytes perturbs oocyte growth and maturation, RNA stability, and the transcriptome. Biol Reprod 2011; 85:575–583

23. Yu J, Deng M, Medvedev S, Yang J, Hecht NB, Schultz RM. Transgenic RNAi-mediated reduction of MSY2 in mouse oocytes results in reduced fertility. Dev Biol 2004; 268:195–206

24. Deng Y, Zhang W, Su D, Yang Y, Ma Y, Zhang H, Zhang S. Some single nucleotide polymorphisms of MSY2 gene might contribute to susceptibility to spermatogenic impairment in idiopathic infertile men. Urology 2008; 71:878–882

25. Bouvet P, Wolffe AP. A role for transcription and FRGY2 in masking maternal mRNA within Xenopus oocytes. Cell 1994; 77:931–941

26. Lau ES, Zhu B, Sun MA, Ngai SM, Ge W. Proteomic analysis of zebrafish folliculogenesis identifies YB-1 (Ybx1/ybx1) as a potential gatekeeping molecule controlling early ovarian folliculogenesis. Biol Reprod 2023; 109:482–497

27. Chen W, Ge W. Ontogenic expression profiles of gonadotropins (*fshb* and *lhb*) and growth hormone (*gh*) during sexual differentiation and puberty onset in female zebrafish. Biol Reprod 2012; 86:73

28. Ge W, Peter RE. Activin-like peptides in somatotrophs and activin stimulation of growth hormone release in goldfish. Gen Comp Endocrinol 1994; 95:213–221

29. Wang Y, Ge W. Gonadotropin regulation of follistatin expression in the cultured ovarian follicle cells of zebrafish, *Danio rerio*. Gen Comp Endocrinol 2003; 134:308–315

30. Zhu B, Pardeshi L, Chen Y, Ge W. Transcriptomic analysis for differentially expressed genes in ovarian follicle activation in the zebrafish. Front Endocrinol (Lausanne) 2018; 9:593. doi:510.3389/fendo.2018.00593

31. Zhou R, Tsang AH, Lau SW, Ge W. Pituitary adenylate cyclase-activating polypeptide (PACAP) and its receptors in the zebrafish ovary: evidence for potentially dual roles of PACAP in controlling final oocyte maturation. Biol Reprod 2011; 85:615–625

32. Pang Y, Ge W. Gonadotropin regulation of activin βA and activin type IIA receptor expression in the ovarian follicle cells of the zebrafish, *Danio rerio*. Mol Cell Endocrinol 2002; 188:195–205

33. Liu KC, Ge W. Differential regulation of gonadotropin receptors (*fshr* and *lhcgr*) by epidermal growth factor (EGF) in the zebrafish ovary. Gen Comp Endocrinol 2013; 181:288–294

34. Chen W, Ge W. Gonad differentiation and puberty onset in the zebrafish: evidence for the dependence of puberty onset on body growth but not age in females. Mol Reprod Dev 2013; 80:384–392

35. Zhai Y, Zhang X, Zhao C, Geng R, Wu K, Yuan M, Ai N, Ge W. Rescue of *bmp15* deficiency in zebrafish by mutation of *inha* reveals mechanisms of BMP15 regulation of folliculogenesis. PLoS Genet 2023; 19:e1010954

36. Zhao C, Zhai Y, Geng R, Wu K, Song W, Ai N, Ge W. Genetic analysis of activin/inhibin β subunits in zebrafish development and reproduction. PLoS Genet 2022; 18:e1010523

37. Chen W, Zhai Y, Zhu B, Wu K, Fan Y, Zhou X, Liu L, Ge W. Loss of growth differentiation factor 9 causes an arrest of early folliculogenesis in zebrafish-A novel insight into its action mechanism. PLoS Genet 2022; 18:e1010318

38. Lu H, Zhao C, Zhu B, Zhang Z, Ge W. Loss of inhibin advances follicle activation and female puberty onset but blocks oocyte maturation in zebrafish. Endocrinology 2020; 161:1–19

39. Hu Z, Ai N, Chen W, Wong QW, Ge W. Leptin and its signaling are not involved in zebrafish puberty onset. Biol Reprod 2022; 106:928–942

40. Budkina KS, Zlobin NE, Kononova SV, Ovchinnikov LP, Babakov AV. Cold Shock Domain Proteins: Structure and Interaction with Nucleic Acids. Biochemistry (Mosc) 2020; 85:S1–S19

41. Evdokimova V, Ruzanov P, Imataka H, Raught B, Svitkin Y, Ovchinnikov LP, Sonenberg N. The major mRNA-associated protein YB-1 is a potent 5’ cap-dependent mRNA stabilizer. EMBO J 2001; 20:5491–5502

42. Wu K, Zhai Y, Qin M, Zhao C, Ai N, He J, Ge W. Genetic evidence for differential functions of *figla* and *nobox* in zebrafish ovarian differentiation and folliculogenesis. Commun Biol 2023; 6:1185

43. Dranow DB, Hu K, Bird AM, Lawry ST, Adams MT, Sanchez A, Amatruda JF, Draper BW. Bmp15 Is an oocyte-produced signal required for maintenance of the adult female sexual phenotype in zebrafish. PLoS Genet 2016; 12:e1006323

44. Ge W. Intrafollicular paracrine communication in the zebrafish ovary: the state of the art of an emerging model for the study of vertebrate folliculogenesis. Mol Cell Endocrinol 2005; 237:1–10

45. Ge W. Paracrine control of fish ovarian follicle development and function. In: Garcia-Ayala A, Penalver JM, Chaves-Pozo E, eds. Recent Advances in Fish Reproduction Biology. Kerala, India: Research Signpost; 2010:141–173.

46. Chung CK, Ge W. Epidermal growth factor differentially regulates activin subunits in the zebrafish ovarian follicle cells via diverse signaling pathways. Mol Cell Endocrinol 2012; 361:133–142

47. Liu KC, Lin SW, Ge W. Differential regulation of gonadotropin receptors (*fshr* and *lhcgr*) by estradiol in the zebrafish ovary involves nuclear estrogen receptors that are likely located on the plasma membrane. Endocrinology 2011; 152:4418–4430

48. Li CW, Ge W. Spatiotemporal expression of bone morphogenetic protein family ligands and receptors in the zebrafish ovary: a potential paracrine-signaling mechanism for oocyte-follicle cell communication. Biol Reprod 2011; 85:977–986

49. Wang Y, Ge W. Involvement of cyclic adenosine 3’,5’-monophosphate in the differential regulation of activin βA and βB expression by gonadotropin in the zebrafish ovarian follicle cells. Endocrinology 2003; 144:491–499

50. Poon SK, So WK, Yu X, Liu L, Ge W. Characterization of inhibin α subunit (*inha*) in the zebrafish: evidence for a potential feedback loop between the pituitary and ovary. Reproduction 2009; 138:709–719

51. Yin Q, Zheng M, Luo Q, Jiang D, Zhang H, Chen C. YB-1 as an oncoprotein: functions, regulation, post-translational modifications, and targeted therapy. Cells 2022; 11:1217. doi:1210.3390/cells11071217

52. Yang F, Cui P, Lu Y, Zhang X. Requirement of the transcription factor YB-1 for maintaining the stemness of cancer stem cells and reverting differentiated cancer cells into cancer stem cells. Stem Cell Res Ther 2019; 10:233

53. Wu J, Lee C, Yokom D, Jiang H, Cheang MC, Yorida E, Turbin D, Berquin IM, Mertens PR, Iftner T, Gilks CB, Dunn SE. Disruption of the Y-box binding protein-1 results in suppression of the epidermal growth factor receptor and HER-2. Cancer Res 2006; 66:4872–4879

54. Harada K, Ogden GR. An overview of the cell cycle arrest protein, p21(WAF1). Oral Oncol 2000; 36:3–7

55. Jung YS, Qian Y, Chen X. Examination of the expanding pathways for the regulation of p21 expression and activity. Cell Signal 2010; 22:1003–1012

56. Niculescu AB, 3rd, Chen X, Smeets M, Hengst L, Prives C, Reed SI. Effects of p21(Cip1/Waf1) at both the G1/S and the G2/M cell cycle transitions: pRb is a critical determinant in blocking DNA replication and in preventing endoreduplication. Mol Cell Biol 1998; 18:629–643

57. Nakanishi M, Adami GR, Robetorye RS, Noda A, Venable SF, Dimitrov D, Pereira-Smith OM, Smith JR. Exit from G0 and entry into the cell cycle of cells expressing p21Sdi1 antisense RNA. Proc Natl Acad Sci USA 1995; 92:4352–4356

58. Deng C, Zhang P, Harper JW, Elledge SJ, Leder P. Mice lacking p21CIP1/WAF1 undergo normal development, but are defective in G1 checkpoint control. Cell 1995; 82:675–684

59. Frye BC, Halfter S, Djudjaj S, Muehlenberg P, Weber S, Raffetseder U, En-Nia A, Knott H, Baron JM, Dooley S, Bernhagen J, Mertens PR. Y-box protein-1 is actively secreted through a non-classical pathway and acts as an extracellular mitogen. EMBO Rep 2009; 10:783–789

60. Pagano C, di Martino O, Ruggiero G, Maria Guarino A, Mueller N, Siauciunaite R, Reischl M, Simon Foulkes N, Vallone D, Calabro V. The tumor-associated YB-1 protein: new player in the circadian control of cell proliferation. Oncotarget 2017; 8:6193–6205

61. Laranjeiro R, Tamai TK, Peyric E, Krusche P, Ott S, Whitmore D. Cyclin-dependent kinase inhibitor p20 controls circadian cell-cycle timing. Proc Natl Acad Sci USA 2013; 110:6835–6840

62. Jurchott K, Bergmann S, Stein U, Walther W, Janz M, Manni I, Piaggio G, Fietze E, Dietel M, Royer HD. YB-1 as a cell cycle-regulated transcription factor facilitating cyclin A and cyclin B1 gene expression. J Biol Chem 2003; 278:27988–27996

